# Domestic dog introgression in Australian dingoes: environmental drivers and evolutionary consequences

**DOI:** 10.64898/2026.03.22.713106

**Authors:** Carolina Osuna-Mascaró, Kylie M. Cairns, Karolina Doan, Alejandro Flores-Manzanero, Bradley J. Nesbitt, Thomas M. Newsome, Małgorzata Pilot

## Abstract

Introgressive hybridization between wild and domestic animals is a widespread phenomenon with important implications for genetic diversity, local adaptation, and conservation management. The causes and consequences of this process are poorly understood. In Australia, hybridization between dingoes and domestic dogs presents a dual conservation challenge, threatening the genetic integrity of dingoes while allowing potential adaptive introgression. To investigate the environmental drivers of this process, we analyzed high-density SNP array data in 390 dingoes and 396 domestic dogs. Dingo populations showed regional genetic structure and were clearly differentiated from domestic dogs. Using local ancestry inference and genome–environment association analyses, we found low levels of dog introgression in dingoes from remote areas in Central and Western Australia, and moderate levels in Eastern and Southern populations. Climatic variables (maximum temperature of the warmest month, mean temperature of the driest quarter) and the Human Footprint Index (reflecting density of human populations and environmental modifications) were significant predictors of introgression. We identified four genomic regions with overrepresented dog ancestry, including a large introgressed block on chromosome 27, which contained an olfactory receptor gene showing signatures of positive selection, suggesting adaptive introgression. In addition, a chromosomal inversion previously described in dogs and absent in dingoes was initially identified as an introgressed block. We also detected eight genomic regions nearly free of dog ancestry, suggesting purifying selection against maladaptive variants. Together, these results highlight the complex interplay between introgression, human influence, and local adaptation in dingoes, offering valuable insights for conserving the evolutionary potential of this apex predator in increasingly modified landscapes.

## 1. Introduction

Hybridization is a natural and widespread phenomenon that has long captivated evolutionary biologists. Early conceptual frameworks provided key insights into its role in speciation, challenging traditional views on how species boundaries are maintained (Dobzhansky, 1937; Mayr, 1942; Stebbins, 1959; Grant and Grant, 1979). This process may involve gene flow between genetically distinct lineages, leading to diverse outcomes such as the extinction or displacement of parental taxa, the fusion of previously divergent taxa, or the formation of new hybrid lineages that may eventually result in speciation (Ellstrand and Elam, 1993; Rieseberg and Wendel, 1993; Mallet, 2007; Abbott et al., 2013; Thomas, 2015; Grant and Grant, 2017). However, introgression, the integration of genetic variation into a recipient population through hybridization followed by backcrossing, constitutes a key mechanism by which alleles move across species boundaries (Aguillon et al., 2022). Advances in genomic technologies have enabled the study of introgression at a genome-wide scale, highlighting its evolutionary relevance and providing deeper insights into its role in shaping biodiversity across diverse taxonomic groups, including fungi (Giraud et al., 2008; Kinnerberg et al., 2023), plants (Mallet, 2007; Stull et al., 2023), fish (Seehausen, 2004; Blumer et al., 2024; Kato et al., 2024), birds (Grant and Grant, 2017; Singhal et al., 2021), and mammals (Leonard et al., 2013; Adavoudi and Pilot, 2021; Tensen and Fisher, 2024).

Among mammals, introgression between canid species is a widespread phenomenon with significant implications for genetic diversity, ecological dynamics, and evolutionary trajectories (Leonard et al., 2013; Adavoudi and Pilot, 2021). This process may be influenced by anthropogenic activities driving hybridization, especially in ecosystems where wild canids like grey wolves (Canis lupus), coyotes (Canis latrans), and golden jackals (Canis aureus) overlap with domestic dogs (Canis familiaris) (Wheeldon et al., 2013; Pilot et al., 2018; McFarlane and Pemberton, 2019; Stefanović et al., 2024). In regions such as North America and Europe, where free-ranging domestic dogs coexist with wild representatives of the genus Canis, hybridization is well-documented and frequently results in the introgression of dog-specific genetic variation into wild species (Leonard et al., 2013). A notable example is melanism in gray wolves in North America, driven by the beta-defensin gene, a melanocortin pathway gene introduced via historical hybridization with domestic dogs (Anderson et al., 2009). This trait has risen to high frequencies under positive selection in forested habitats, exemplifying adaptive introgression (Anderson et al., 2009). However, hybridization and subsequent introgression may pose risks to the genetic and phenotypic distinctiveness of wild canids, challenging conservation efforts aimed at preserving unique ecological roles and lineage integrity (Hohenlohe et al., 2021).

Wild canids increasingly occupy habitats highly modified by humans and with access to human food waste (Kuijper et al., 2016). In such habitats, wild canids are likely to shift, at least partially, from hunting wild prey to livestock depredation and/or scavenging anthropogenic food (Newsome et al., 2015a). Because evolutionary divergence in canids may be strongly influenced by differences in diet composition (Pilot et al., 2006), dietary shifts may trigger a contemporary domestication process (Newsome et al., 2017). Hybridization with free-ranging dogs may accelerate this process by enabling wild canids to rapidly acquire adaptations to the niche of human commensal (Pilot et al., 2021). While dog-derived traits may be maladaptive in natural habitats, they can be advantageous in human-dominated landscapes. Thus, introgression may facilitate adaptation to anthropogenic habitats but simultaneously shift the ecological niche, with potentially negative ecosystem-level consequences (vonHoldt et al., 2018).

The dingo presents a unique case within canid hybridization systems. Present in Australia for at least 5,000 years (Fillios and Taçon, 2016), dingoes have assumed the role of apex predators, with varying effects on both co-occurring predators and prey (Glen et al., 2007; Letnic and Koch, 2010; Newsome et al., 2015b; Doherty et al., 2019; Castle et al., 2023). The dingo genome displays patterns of natural selection distinct from domestic dogs, reflecting its adaptation to the apex predator niche (Zhang et al., 2020). Dingoes exhibit social and behavioral traits similar to other wild representatives of the genus *Canis*, including pack structures led by a dominant pair, seasonal breeding and the cooperative hunting of large prey like kangaroos and wallabies (Corbett, 1995; Glen et al., 2007; Pollock et al., 2022). Yet, this ecological function is potentially threatened by extirpation, human-wildlife conflict, and hybridization with domestic dogs (Newsome et al., 2015b). For the latter, the arrival of domestic dogs with European origin created new opportunities for interbreeding, and concerns about the need to preserve the genetic integrity of dingo populations are often raised (Stephens et al., 2015; Cairns et al., 2017, 2020).

Genetic studies have highlighted the distinct evolutionary history of dingoes relative to domestic dogs, reflecting dingoes’ ancient origins and genetic uniqueness (Cairns et al., 2017; Cairns et al., 2021; Souilmi et al., 2024). Numerous studies using microsatellites have documented the genetic impact of dingoes interbreeding with dogs, suggesting it may erode dingo genetic integrity in geographic regions characterized by high densities of dogs via genetic swamping (Wilton, 2001; Elledge et al., 2006; Glen et al., 2007; Stephens et al., 2015, 2022; Cairns et al., 2021). This situation has led to complex management challenges (Boronyak et al., 2023). However, recent SNP-based genomic studies suggest that the extent of contemporary interbreeding between dingoes and dogs has been overstated (Cairns et al., 2023; Weeks et al., 2024). Whole genome sequence analysis with an ancient DNA baseline identified 9.7 to 22.5% introgressed European dog ancestry persisting in dingoes from southeast Australia, while minimal dog ancestry was detected in northwest dingoes (Scarsbrook et al., 2025). Consistently, genetic surveys of free-living canines in Australia indicate that domestic dogs and first-generation hybrids are rare (<1%), and genome-wide admixture analyses reveal a bimodal ancestry distribution dominated by either pure dingoes and dingo–dog backcrosses or pure domestic dogs, with minimal numbers of first-generation hybrids (Cairns et al., 2023). A similar pattern is observed in the case of wolf-dog hybridization across Eurasia (Pilot et al., 2021; Lobo et al., 2025; Sarabia et al., 2025; Battilani et al., 2025). Additionally, studies on the skull shape and the ecosystem impacts of dingoes in different parts of their range suggest that introgression from domestic dogs have had a limited effect on the dingo’s ecological role (Parr et al., 2016; Crowther et al., 2021). Yet, several key questions remain. First, discrepancies between previous studies highlight uncertainty about the true extent of introgression, with concerns that estimates may be biased by underlying population structure and detection of historical introgression. Second, while human activities such as domestic dog ownership and lethal control have been suggested as potential drivers of increased hybridization, the specific environmental factors influencing admixture rates are not well understood. Finally, the evolutionary consequences of introgression for dingoes, especially the possibility of adaptive introgression, remain largely unexplored.

To address these knowledge gaps, we combined landscape genomics, local ancestry inference, and introgression analyses, using DNA samples collected from dingoes and European and Australian domestic dogs. We applied high-resolution local ancestry methods to improve detection accuracy, explicitly accounted for population structure, and explored how environmental factors shape spatial patterns of introgression between dingoes and dogs. We further examined introgressed genomic regions for signatures of selection, providing insights into the evolutionary impact of dog ancestry on dingoes and informing conservation efforts.

## 2. Material and Methods

### 2.1 Sample collection and SNP genotyping

Tissue samples were collected from wild dingoes across Australia, primarily opportunistically following lethal management activities or retrieved as roadkill by private citizens. We also collected blood or buccal swab samples from Australian domestic pet dogs, selecting commonly occurring working, herding, and mixed breeds to capture the diversity most likely to interact with regional dingo populations. DNA was extracted from blood, tissue, or buccal samples using Qiagen DNeasy Blood and Tissue kits (Qiagen Sciences, Germantown, USA). Extracted DNA was genotyped at the Ramaciotti Centre for Genomics (University of New South Wales, Randwick, Australia) on the Axiom Canine HD Genotyping array (Thermo Fisher Scientific Inc., Waltham, USA). In addition, we incorporated previously published genotype data from Cairns et al. (2023), generated on the same Axiom Canine HD array, to augment our representation of wild dingo diversity. This research complies with applicable laws on sampling from natural populations and animal experimentation, including the ARRIVE guidelines (Du Sert et al., 2020).

The combined dataset analyzed in this study comprises genotypic data from 300,761 SNPs following rigorous QC and 170,465 SNPs following further LD-pruning in PLINK v1.9 (Purcell et al., 2007). Filtering steps included removing individuals with more than 10% of missing data (option --mind 0.1) and excluding markers based on missingness (--geno 0.1), minor-allele frequency (--maf 0.01), and LD (--indep-pairwise 50 10 0.5). Finally, individuals with relatedness up to second-degree were removed using --king-cutoff 0.0885 option in PLINK v2.0. To ensure balanced representation, we included equal numbers of dingoes and European dogs (both purebred and free-ranging). Our dataset contained 390 dingoes sampled from multiple regions across Australia (Big Desert, Central, East, North, South, West, and captive populations), along with 54 domestic dogs representing Australian breeds (including 33 purebred dogs and 21 mixed-breed dogs), 116 European free-ranging dogs, 226 European purebred dogs (i.e. dog breeds of European origin sampled in the United States), and two F1 dog–dingo hybrids (Table S1). Our sampling strategy for dingoes was population-focused, with multiple individuals collected from specific localities to facilitate analyses of the influence of local environmental factors and human footprint on introgression dynamics. Australian breeds are breeds developed in Australia based on domestic dogs brought from Europe. Because we sampled owned dogs that do not range freely and represent breeds whose reproduction is managed by humans, these individuals are unlikely to represent populations that routinely hybridise with dingoes. However, occasional reports exist of owned farm dogs breeding with wild dingoes. Our sampled individuals therefore primarily represent managed lineages of domestic dogs introduced from Europe, rather than free-ranging or feral dog populations directly involved in contemporary hybridisation events, while still capturing the ancestral genetic background shared with many rural Australian dogs. European dogs represent the parental population for the Australian dogs and were included so that the broader European gene pool is represented, to account for potential unsampled dog lineages that were introduced to Australia and interbred with dingoes. European pure-breeds were drawn from Morrill et al. (2022), excluding any breeds of non-European origin (e.g. Afghan Hound, Chow Chow). European free-ranging dogs were sampled from across Eastern Europe and include samples previously used in Spatola et al. (2023).

### 2.2 Ancestry Analysis

To investigate how introgression patterns vary across Australia and to identify genomic regions associated with introgression, we employed a combination of global ancestry analyses (which provide an overall estimate of each individual’s ancestral composition across the entire genome) and local ancestry analyses (which detect chromosome-level ancestry variation). Local ancestry inference can reveal signatures of older admixture events that have since become pervasive across the genome, enabling the detection of historical introgression that might be overlooked by global methods (Sankararaman et al., 2014). By integrating both approaches, we captured broad admixture patterns as well as discrete regions of introgression.

First, we assessed the genome-wide genetic structure using principal component analysis (PCA) (Patterson et al., 2006) and ADMIXTURE (Alexander and Lange, 2011) to identify distinct populations of dingoes across Australia that may differ in introgression patterns. These analyses were carried out for the entire dataset and for the dingo dataset alone (see Supplementary Material for details).

To examine local ancestry and introgression signals at the chromosome-level resolution, we employed two complementary methods: LAMP-LD software v2.4 (Sankararaman et al., 2008) and ELAI software (Guan, 2014), both performed on a dataset without LD-pruning (300,761 SNPs) to preserve haplotype information and maximize the resolution of local ancestry tracts. LAMP-LD was selected for its ability to infer local ancestry without the prior designation of "pure" reference populations, making it particularly useful for providing initial estimates of local ancestry in datasets with complex population structures. We divided the dataset into chromosome-specific files using PLINK v1.09 (Purcell et al., 2007). Custom bash scripts were used to configure LAMP-LD parameters for each of the 38 autosomal chromosomes (see Data Accessibility section).

ELAI was then applied to refine local ancestry estimates, leveraging its flexibility in handling dense SNP data and its ability to model complex population histories. The dataset was divided into reference populations (individuals with high dingo and domestic dog ancestry, respectively, identified based on the LAMP-LD results) and admixed populations, which included admixed dingoes, known hybrids and Australian mixed-breed dogs. Genotype data were then converted from PLINK to the ’bimbam’ format required for ELAI input files. ELAI was executed with parameters optimized for local ancestry inference: -mg was set to 10 to specify the maximum generations since admixture, and -C was set to 2 to assume two ancestral populations (dingoes and domestic dogs). The parameter -c was set to 10 to increase the flexibility of the hidden Markov model in capturing complex local patterns of ancestry across haplotypes, which enhances the model’s ability to detect fine-scale introgression signals. Additionally, -R was set to 45 to optimize the number of EM iterations and ensure convergence. The results were summarized to visualize the mean proportions of dingo and dog ancestry across the genome using ggplot2 in R (Wickham and Wickham, 2016).

Finally, we used the GHap package (Utsunomiya et al., 2016) to investigate the distribution of extended haplotype blocks across the genome, providing complementary information to the inferences obtained with LAMP-LD and ELAI. GHap focuses on identifying and analyzing extended haplotype structures, enabling the detection of broader patterns of recombination, shared ancestry, and structural signals of introgression that SNP-based local ancestry methods may miss. Prior to the analysis, all genotypes were phased using Beagle v5.4 (Browning et al., 2021), with the burn-in parameter set to 10 and iterations to 1000, to ensure accurate haplotype reconstruction. We applied GHap to the LD-unpruned dataset, retaining all loci to fully capture LD-based haplotype structure. An unsupervised GHap analysis was performed using the elbow method to determine the optimal number of clusters (K), which was found to be K = 2, corresponding to dingo and dog ancestral populations. Individuals with ancestral purity exceeding 90% were then classified as non-admixed dingoes or dogs. This information was used in a supervised GHap analysis to examine introgression events and further refine estimates of genetic structure. The results were visualized with karyoplots and Manhattan plots in R, highlighting the genomic distribution of ancestral contributions and identifying potential introgression events.

### 2.3 Detection and Characterization of Introgressed Blocks

Potential adaptive introgression in the dingo population was investigated by analyzing regions of elevated dog ancestry across the genome to identify candidate genes under adaptive introgression. Ancestral allele dosage data obtained from ELAI were analyzed for all chromosomes (1-38). SNP information files (“.snpinfo.txt”) and dosage files (“.ps21.txt”) were processed to ensure consistent data dimensions, and dog introgression rates were calculated by averaging dog dosage values across all dingoes and rescaling the resulting proportions to the [0, 1] range. Regions with elevated dog ancestry were identified using a conservative, chromosome-specific threshold, defined as SNPs with ancestry values exceeding three standard deviations above the mean for each chromosome. This threshold was used as a descriptive criterion to highlight genomic regions showing unusually high ancestry relative to the chromosomal background, rather than as a formal SNP-wise hypothesis test, as local ancestry estimates are highly autocorrelated along chromosomes. We chose this chromosome-specific approach, following Pilot et al. (2021), rather than a single genome-wide cutoff, because each chromosome acts as an independent recombination unit, and ancestry blocks are expected to vary in size and distribution across chromosomes. Automated R scripts facilitated the visualization and comparison of the elevated-ancestry segments across all chromosomes. Conversely, “ancestry deserts” (regions of exceptionally low dog ancestry; Kim et al., 2018) were identified following Sankararaman et al. (2014) as runs of ≥10 consecutive SNPs with <0.1% dog ancestry.

To support the interpretation of the chromosome 9 candidate region (see Results), we additionally estimated Weir and Cockerham’s F_ST_ in sliding windows along chromosome 9. F_ST_ was calculated in 50 kb non-overlapping windows between dingoes and domestic dogs using VCF-based genotype data.

To evaluate whether ancestry deserts harbor an excess of potentially deleterious dog-associated variants, we annotated variants within these regions using Ensembl Variant Effect Predictor (VEP; McLaren et al., 2016). We focused on variants showing strong allele-frequency differences between dogs and dingoes, retaining sites segregating in dogs but rare or absent in dingoes (AF_dogs ≥ 0.05 and AF_dingoes ≤ 0.01). As a sensitivity analysis, we repeated this using a more permissive threshold (AF_dogs ≥ 0.02; AF_dingoes ≤ 0.01). We summarized predicted functional consequences and screened for missense and loss-of-function annotations (including SIFT-deleterious missense and stop-gained, frameshift, and essential splice-site variants).

Candidate introgression blocks were identified across different chromosomes using SNP array data, and their genomic coordinates were mapped to the Canis familiaris reference genome (canFam6; assembly ID: GCF_000002285.5) using the UCSC Genome Browser, which served as a common coordinate and annotation framework. SNP positions from the Axiom Canine HD array, originally defined on earlier canine genome assemblies, were converted to canFam6 coordinates using established UCSC liftover mappings prior to downstream analyses. Genes located within these regions were identified based on canFam6 annotations, and their predicted functions were manually verified using GeneCards (Safran et al., 2010). Chromosomes displaying prominent or consistent introgression patterns, along with those containing isolated peaks of interest, were selected for further analyses. To assess patterns of molecular evolution within introgressed regions, BED files defining introgressed blocks were generated. Coding sequences corresponding to genes located within these regions were extracted from the dingo reference genome (Canis lupus dingo; GCA_003254725.2) using BEDTools v2.29.2 (Quinlan, 2014). Homologous coding sequences from the grey wolf reference genome (Canis lupus; GCA_905319855.2) were included as an outgroup for comparative analyses. Introgressed regions themselves were defined exclusively based on SNP array data.

Extracted sequences were converted to FASTA format, and custom Python scripts were employed to correct frameshifts (see Data Accessibility section). Multiple sequences were aligned using MAFFT v7.450 (Katoh and Standley, 2014) to ensure consistency across analyses. To evaluate potential selection pressures acting on introgressed regions, dN/dS ratios were calculated by comparing aligned coding sequences from the canFam6 reference genome and domestic dogs, with the grey wolf (Canis lupus) included as an outgroup. The Ka/Ks ratio, implemented in the R packages ape v5.8 (Paradis et al., 2018) and seqinr v4.2-36 (Charif et al., 2023), was used to further evaluate selection pressures. Statistical significance was assessed using z-tests, focusing on genes with dN/dS ratios that significantly deviated from neutrality (dN/dS ≠ 1; Kryazhimskiy and Plotkin, 2008), thereby identifying candidate genes potentially under positive selection.

To assess the extent of linkage disequilibrium (LD) within introgressed blocks, we computed pairwise LD values (R²) using PLINK v1.9. LD calculations were performed separately for each chromosome. R² values were extracted from the resulting LD matrices, and their distribution was summarized for each chromosome to evaluate patterns of haplotype structure and recombination. Histograms and summary statistics were generated in R to visualize the spread of LD values, allowing us to determine whether introgressed regions exhibit extended haplotype blocks, indicative of reduced recombination.

### 2.4 Introgression analysis based on D-statistics

Variant calling was performed jointly across all canid genomes analysed in the study, including dingoes, domestic dogs, New Guinea Singing Dogs (NGSD), grey wolves, and the golden jackal, generating a single multi-sample variant call set. Raw variants were called using bcftools mpileup and bcftools call (Narasimhan et al., 2016) and filtered to retain high-confidence biallelic SNPs (QUAL ≥ 30, DP ≥ 10). This joint VCF constituted the basis for all subsequent phylogenetic and introgression analyses.

To reconstruct maximum likelihood phylogenetic trees for dingoes and domestic dogs, we used IQ-TREE v3 (Nguyen et al., 2015; Minh et al., 2020). Genome-wide SNP data included in the joint variant call set were available for 390 dingoes distributed across regional populations and 400 domestic dogs (200 European purebred dogs, 42 Australian purebred dogs, 19 Australian mixed-breed dogs, and 139 European free-ranging dogs), as well as 44 grey wolves (*Canis lupus*), one golden jackal (*Canis aureus*) from Stefanovic et al. (2024), and two New Guinea Singing Dogs (NGSD; NCBI accessions SRR7107989 and SRR7107990). The golden jackal was used as the outgroup. SNPs for the two NGSD genomes were extracted from the same joint variant call set using identical filtering criteria, ensuring full comparability across taxa. Individuals identified as recent backcrosses (N = 3) based on local ancestry analyses (ELAI) were excluded prior to phylogenetic reconstruction. Model selection was performed using the ModelFinder Plus option based on the Bayesian Information Criterion (BIC), and branch support was assessed using ultrafast bootstrapping with 1,000 replicates. Phylogenetic trees were visualized using FigTree v1.4.4 (Rambaut, 2009).

To investigate signatures of introgression between dingoes and dogs, we estimated D statistics (ABBA–BABA test) and the f_4_-ratio using Dtrios from Dsuite v0.1 (Malinsky et al., 2021) across all autosomal chromosomes. Analyses were performed on the global dataset as well as separately for each regional dingo population. We used chromosome-specific VCF files and the previously inferred maximum likelihood phylogeny to define population relationships. Individuals identified as recent backcrosses based on local ancestry (ELAI) were excluded from the analysis. Briefly, the ABBA–BABA test examines introgression within a four-taxon topology of (((P1, P2), P3), O), where O is the outgroup. The D statistic compares the frequency of ABBA (derived allele shared between P2 and P3) and BABA (derived allele shared between P1 and P3) site patterns. Under the null hypothesis of no gene flow between P3 and either P1 or P2, ABBA and BABA patterns are expected at equal frequencies; significant deviations from this expectation indicate excess allele sharing.

For our analyses, we focused on trios involving dingoes and domestic dogs, structured as ((NGSD, Dingo), Dog), using the golden jackal as the outgroup. This configuration allows detection of excess shared derived alleles between dingoes and domestic dogs beyond what is expected from shared ancestry with NGSD, which represents a closely related lineage (Surbakti et al., 2020; Souilmi et al., 2024). We acknowledge that the captive NGSD population derives from a limited number of founders and has experienced substantial inbreeding and genetic drift. However, despite this demographic history, NGSD remains the closest available non-dingo lineage and provides a relevant phylogenetic reference for testing excess allele sharing. Although NGSD may also experience introgression from domestic dogs of European origin, such gene flow would be independent from the introgression in dingoes and its extent in captive NGSD individuals analysed here is expected to be limited (Scarsbrook et al., 2025). Therefore, excess allele sharing detected between dingoes and domestic dogs relative to NGSD is unlikely to be explained by shared ancestral polymorphism alone.

We also applied this approach to the introgressed block identified on chromosome 27. We extracted SNPs corresponding to this region using bcftools (Narasimhan et al., 2016) and re-ran Dtrios to test for excess allele sharing at these loci. This refined analysis assessed whether excess allele sharing was elevated locally relative to genome-wide expectations, providing support for localized dog-derived ancestry in the dingo genome.

To explicitly address the limitations of D-statistics in distinguishing introgression from incomplete lineage sorting at local genomic scales (Martin et al., 2015), we additionally quantified localized introgression across the chromosome 27 candidate block using the f_d statistic implemented in Dsuite (Dinvestigate). f_d was estimated in sliding windows across the block to assess whether excess allele sharing with domestic dogs was concentrated within this region, providing a more robust measure of localized introgression independent of genome-wide background processes.

### 2.5 Analysis of population-level variation in introgression using Bayesian genomic clines

Because population-level differences in introgression may not be fully captured by genome-wide or predictive approaches, we additionally applied Bayesian genomic cline analyses to characterize heterogeneity in introgression across loci within each population implemented in the R package bgc-hm (Gompert et al., 2024). This approach models locus-specific deviations in ancestry transitions along a hybrid index while explicitly accounting for genome-wide and population-level variance. Analyses were conducted separately for each dingo population (Big Desert, Central, East, South, West, and captive), using a shared set of ancestry-informative SNPs derived from allele frequency differences between dingoes and domestic dogs. For each population, individual hybrid indices were first estimated under a diploid genotype model using the function est_hi. These estimates were then used as input for hierarchical genomic cline models (est_genocl), which jointly estimate cline centers (α) and cline slopes (β) across loci while allowing for population-specific dispersion parameters. To summarize introgression heterogeneity within populations, we extracted posterior distributions of the dispersion of cline centers (SDc) and cline slopes (SDv). Higher SDc or SDv values indicate greater locus-specific heterogeneity in introgression, whereas lower values reflect more homogeneous ancestry transitions across the genome. All models were run for 3,000 MCMC iterations using default priors, and convergence was assessed visually.

### 2.6 Genetic and Environmental Drivers of Dingo-Dog Introgression Patterns

To analyze the potential role of environmental variables in shaping dingo-dog introgression patterns, we first compiled topographic, environmental and landscape data potentially driving these patterns (Table S2). Topographic (elevation) and environmental (19 bioclimatic variables) data were obtained from the WorldClim database (Hijmans et al., 2005). These variables represent annual trends (e.g., mean annual temperature, annual precipitation), seasonality (e.g., temperature seasonality, precipitation seasonality), and extreme or limiting environmental factors (e.g., temperature of the coldest and warmest months, precipitation of the driest and wettest quarters). The WorldClim data were based on averages for the years 1970–2000. Landscape data were obtained from the Australian Bureau of Agricultural and Resource Economics and Sciences data package (ABARES, 2022) and comprised land use and vegetation cover for Australia from 2010-2011 and 2015-2016. This included land use change, trees, shrub and croplands cover. Additionally, we incorporated regional human population data and dingo barrier fence data obtained from the Australian Bureau of Statistics (ABS, 2023). Human footprint index data from the Global Human Settlement Layer (GHSL; Schiavina et al., 2022) was also included and used for calculating the distance to the nearest human settlement. We set all the variables at 30 seconds of the geographic coordinate system (∼1 km²) of spatial resolution and obtained the environmental data for each sampling location. Multicollinearity among predictors was addressed by calculating pairwise correlations using the “pairs.panels” function from the psych v2.1.9 R package (Revelle and Revelle, 2015), excluding variables with Pearson’s correlation values of *|r|* > 0.70.

We then evaluated associations between genetic variation, local adaptation, and environmental factors by testing for isolation by distance (IBD; Wright, 1943) and isolation by environment (IBE). IBE is observed when positive correlation occurs between genetic and environmental distances, independent of the effect of geographic distance (Wang and Bradburd, 2014). Pairwise genetic distances (Nei’s D) were estimated in R using the package hierfstat v0.5-7 (Goudet, 2005) with the "genet.dist" function and compared to geographic and environmental distances using Mantel tests. Geographic distances were calculated as Haversine distances with the “distm” function from the geosphere v1.5–14 R package (Hijmans et al., 2017), while environmental distances were computed as Euclidean distances using the “dist” function from the stats v3.3.1 R package (R Core Team, 2013). Mantel tests were performed using Spearman correlation with 9999 permutations in the vegan v2.5–7 R package (Oksanen et al., 2013).

We conducted partial Redundancy Analysis (pRDA; Capblancq and Forester, 2021) to partition the variance in genetic variation explained by climate, geography, and population structure. The response variable consisted of individual genotypes (coded as allele counts: 0/1/2), while explanatory variables were grouped into three sets: (1) climate-related environmental variables (Table S2a), (2) proxies of genetic structure (population scores along PC1 and PC2), and (3) geographic coordinates (latitude and longitude). The full pRDA model was fitted using all explanatory variables simultaneously and tested for overall significance. A stepwise procedure (ordiR2step) was applied only as an exploratory analysis to assess the relative contribution of individual predictors, with significance determined by p < 0.01 and adjusted R² values. Three separate pRDA models were performed to evaluate the independent contributions of each set of variables: (1) an environment-only model conditioned on geography and population structure, (2) a population structure-only model conditioned on geography and environmental variables, and (3) a geography-only model conditioned on population structure and environmental variables. This approach disentangled confounding effects and allowed us to assess the relative contributions of each factor. This framework allows environmental associations to be interpreted conservatively by explicitly accounting for spatial structure and shared ancestry, which are major sources of confounding in genotype–environment association analyses.

To investigate potential signals of local adaptation, we performed a second pRDA specifically designed to detect genetic associations with environmental predictors. This analysis used a response matrix of genotypes, with environmental predictors as explanatory variables and population structure (PC1 and PC2) and geography (latitude and longitude) as conditioning variables. The significance of RDA axes and individual environmental predictors was tested using permutation tests (n = 999) with the anova.cca function in vegan. Outlier loci potentially under selection were identified based on SNP loadings along the first three constrained axes, applying a stringent filtering threshold of 3.5 standard deviations (p < 0.0005) (Forester et al., 2018). We manually checked for duplicate candidate loci associated with multiple RDA axes and used Pearson’s correlation (r) to identify the strongest environmental predictor for each outlier locus. This step aimed to detect loci potentially involved in local adaptation, linking genetic variation to environmental drivers. As with other multivariate GEA approaches, results are expected to depend on the set of environmental predictors considered; however, the pRDA framework is well suited to identify robust associations with major climatic gradients while controlling for non-environmental structure.

Random Forest (RF) analyses were used to investigate environmental drivers of variation in introgression across dingo populations. Dog ancestry proportions were estimated at the individual level using ELAI (see above) and used as a continuous response variable in all RF models. We initially considered the full set of environmental and anthropogenic predictors described above, including 19 bioclimatic variables, elevation, land-cover variables, and human footprint metrics (26 predictors in total). To reduce redundancy among predictors, we filtered variables based on pairwise correlations, iteratively removing highly correlated predictors (|r| > 0.8). This procedure resulted in a reduced but still comprehensive set of 16 predictors. No variable selection based on the RDA results was applied, and RF analyses were therefore conducted independently of the pRDA. RF regression models were fitted using the randomForest R package (Liaw and Wiener, 2002), with 1,000 trees and default settings for the number of variables considered at each split. Variable importance was quantified using permutation-based importance measures (%IncMSE), which reflect the decrease in predictive accuracy when each predictor is randomly permuted. Model performance was first evaluated using out-of-bag (OOB) predictions. To further assess model generalizability and reduce the risk of overfitting, we implemented a leave-one-population-out cross-validation (LOPO) approach. In this framework, each population was sequentially excluded from model training and used exclusively for testing, providing an explicit out-of-sample evaluation of predictive performance. Model accuracy was quantified using mean squared error (MSE) and the coefficient of determination (R²).

## 3. Results

### 3.1 Population Structure and Admixture Patterns

The PCA of both dingo and dog populations of European origin revealed clear genetic structuring, with dingoes forming a distinct cluster separate from domestic dogs. Free-ranging and purebred dogs exhibited genetic differentiation but were more closely related to each other than to dingoes (Figure 1a). ADMIXTURE analyses (K = 12, Figure S1) supported the PCA results, showing that dingoes and domestic dogs formed separate clusters (Table S3a). ADMIXTURE estimated the average dog admixture in dingoes at 0.09 and dingo admixture in Australian mixed-breed and pure-breed dogs at 0.04 and 0.02, respectively. Dingo admixture in European pure-breed dogs and European free-ranging dogs was estimated at 0.01 and 0.03, respectively, which provides an estimate of a false positive rate.

**Figure 1.**
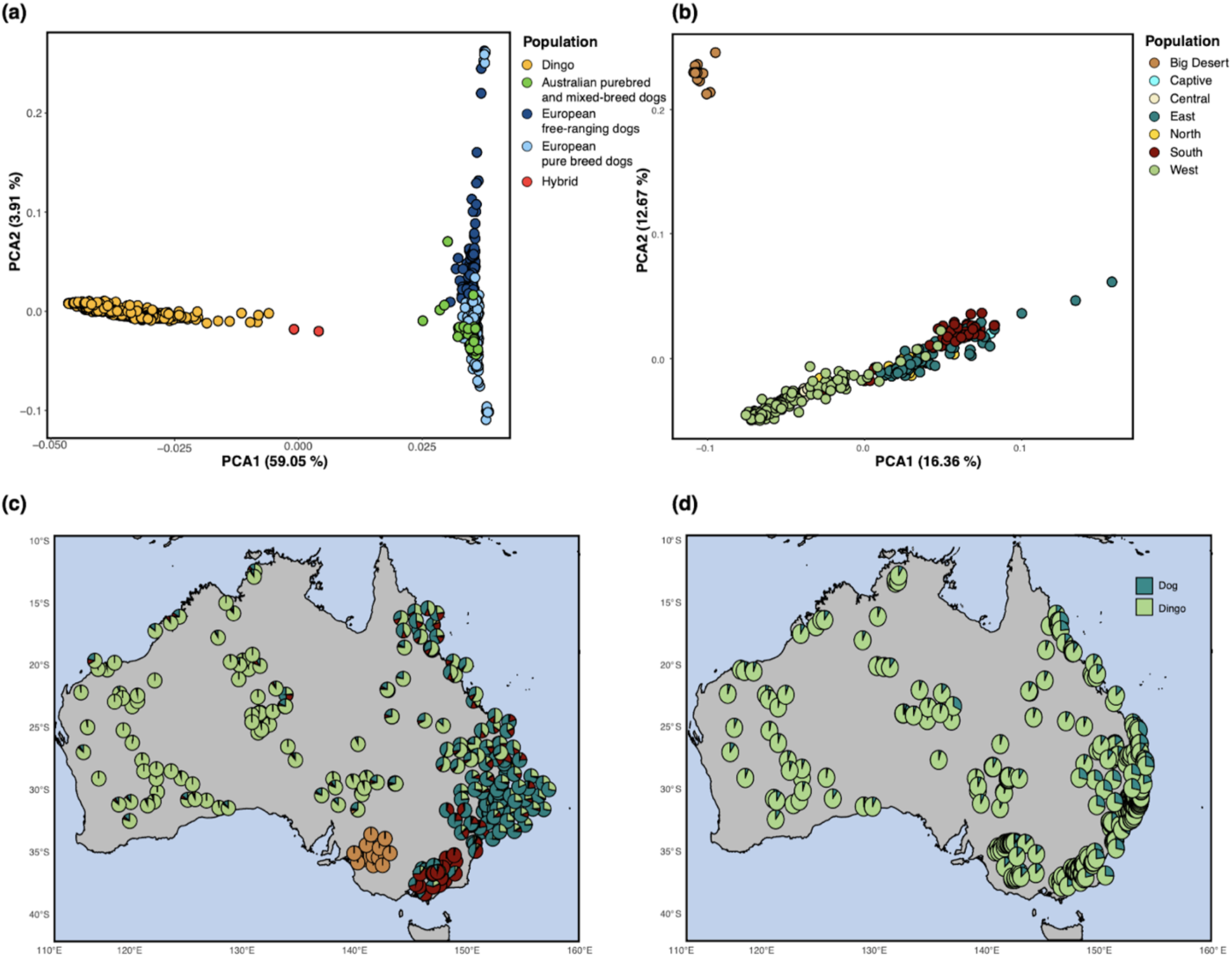
Population structure and genetic differentiation of dingoes and domestic dogs across Australia. **(a)** Principal Component Analysis (PCA) including dingoes, domestic dogs, and putative hybrids. Dingoes are shown in yellow, Australian domestic dogs (purebred and mixed-breed) in green, European free-ranging dogs in dark blue, European purebred dogs in light blue, and hybrids in red. The PCA highlights the strong genetic differentiation between dingoes and domestic dogs, as well as individuals with intermediate genotypes. The percentage of variance explained by each axis is indicated. **(b)** PCA of dingo populations only, colored by geographic group (Big Desert, Captive, Central, East, North, South, and West), showing fine-scale population structure within Australia. **(c)** Geographic distribution of dingo populations across Australia. Pie charts at each sampling location represent the proportion of ancestry from the four genetic clusters (K = 4) inferred by ADMIXTURE (see Figure S2). **(d)** Geographic distribution of local ancestry inferred with ELAI. Pie charts represent the estimated proportion of dingo (green) and domestic dog (blue) ancestry for each individual, illustrating the spatial heterogeneity of introgression across Australia.

When focusing exclusively on the dingo population, the PCA revealed significant genetic differentiation among subpopulations, primarily aligned with geographic regions. Dingoes from the Big Desert and Western Desert formed distinct genetic clusters, while subpopulations from Northern, Southern, and Eastern Australia showed varying degrees of overlap, reflecting regional differentiation and possible gene flow (Figure 1b). ADMIXTURE analyses (Figure 1c, Figure S2) corroborated these patterns, highlighting the genetic isolation of desert populations and shared ancestry among Northern and Eastern subpopulations.

### 3.2 Local Ancestry and Introgression Analysis

#### 3.2.1. Local Ancestry Patterns

The combined results from LAMP-LD, ELAI, and GHap analyses revealed clear patterns of dog introgression in dingoes across Australia (Supplementary Table S3). Both LAMP and ELAI estimated the average dog admixture proportion at 0.15 (for ELAI, this estimate was achieved by assuming that all individuals included in the reference population as pure dingoes had dog admixture equal 0), while GHap estimated the admixture proportion at 0.12. All methods identified significant population-level differences in introgression levels, showing that dingoes from the Central and Western region exhibit markedly lower proportions of dog ancestry compared to those from other regions, as well as the captive population, while the highest admixture occurs in the Eastern Region (Figure 1d, Figure S3; Table 1; Table S3b). All three methods estimated dingo admixture in mixed-breed and pure-breed Australian dogs at less than 0.02 and less than 0.01, respectively, implying very limited introgression from dingoes to dogs (Table S3). In European dogs, the estimate of dingo admixture was negligible, showing that the false positive rate (i.e. detection of false admixture) is low for all three methods.

**Table 1.** Sample size (N) and mean proportion of dog ancestry in dingoes, hybrids and reference dog groups, as estimated by four analytical methods: ADMIXTURE (global ancestry), GHap (haplotype-block sharing), LAMP (LD-based local ancestry) and ELAI (two-layer local ancestry). Values represent the average fraction of dog-derived genome across individuals in each population.

| **Population** | **N** | **ADMIXTURE** | **GHap** | **LAMP** | **ELAI** |
| --- | --- | --- | --- | --- | --- |
| **All dingoes** | 390 | 0.091 | 0.123 | 0.151 | 0.151 |
| **Central** | 36 | 0.006 | 0.030 | 0.041 | 0.031 |
| **West** | 109 | 0.037 | 0.061 | 0.079 | 0.075 |
| **Big Desert** | 15 | 0.016 | 0.110 | 0.127 | 0.130 |
| **North** | 49 | 0.116 | 0.149 | 0.181 | 0.186 |
| **South** | 66 | 0.132 | 0.162 | 0.203 | 0.197 |
| **East** | 79 | 0.166 | 0.183 | 0.224 | 0.228 |
| **Captive** | 36 | 0.102 | 0.169 | 0.190 | 0.198 |
| **Dingo-dog F1 hybrid** | 2 | 0.557 | 0.568 | 0.588 | 0.597 |
| **Australian mixed-breed dogs** | 21 | 0.956 | 0.985 | 0.987 | 0.989 |
| **Australian pure-bred dogs** | 33 | 0.980 | 0.993 | 0.993 | 0.998 |
| **European pure-bred dogs** | 226 | 0.989 | 0.999 | 0.998 | 1 |
| **European free-ranging dogs** | 116 | 0.968 | 0.999 | 0.998 | 1 |

All three methods of local ancestry analyses highlighted chromosomes 9 and 27 as genomic regions with particularly elevated dog ancestry (Figure 2), especially in Eastern and Southern populations (Table S3b-d). A comparison of global versus local ancestry estimates revealed that the three local methods (LAMP-LD, ELAI and GHap) produced highly consistent results (mean difference ≈ 1% per individual), whereas global ancestry proportions from ADMIXTURE differed by 6-7% on average from the local ancestry estimates (Table 1; Figure S4). Comparison of ancestry estimates across genomic scales shows that both the mean and variance of dog ancestry increase markedly on chromosomes 9 and 27 relative to genome-wide estimates, with highly concordant patterns between LAMP and ELAI (Figure 2b).

**Figure 2.**
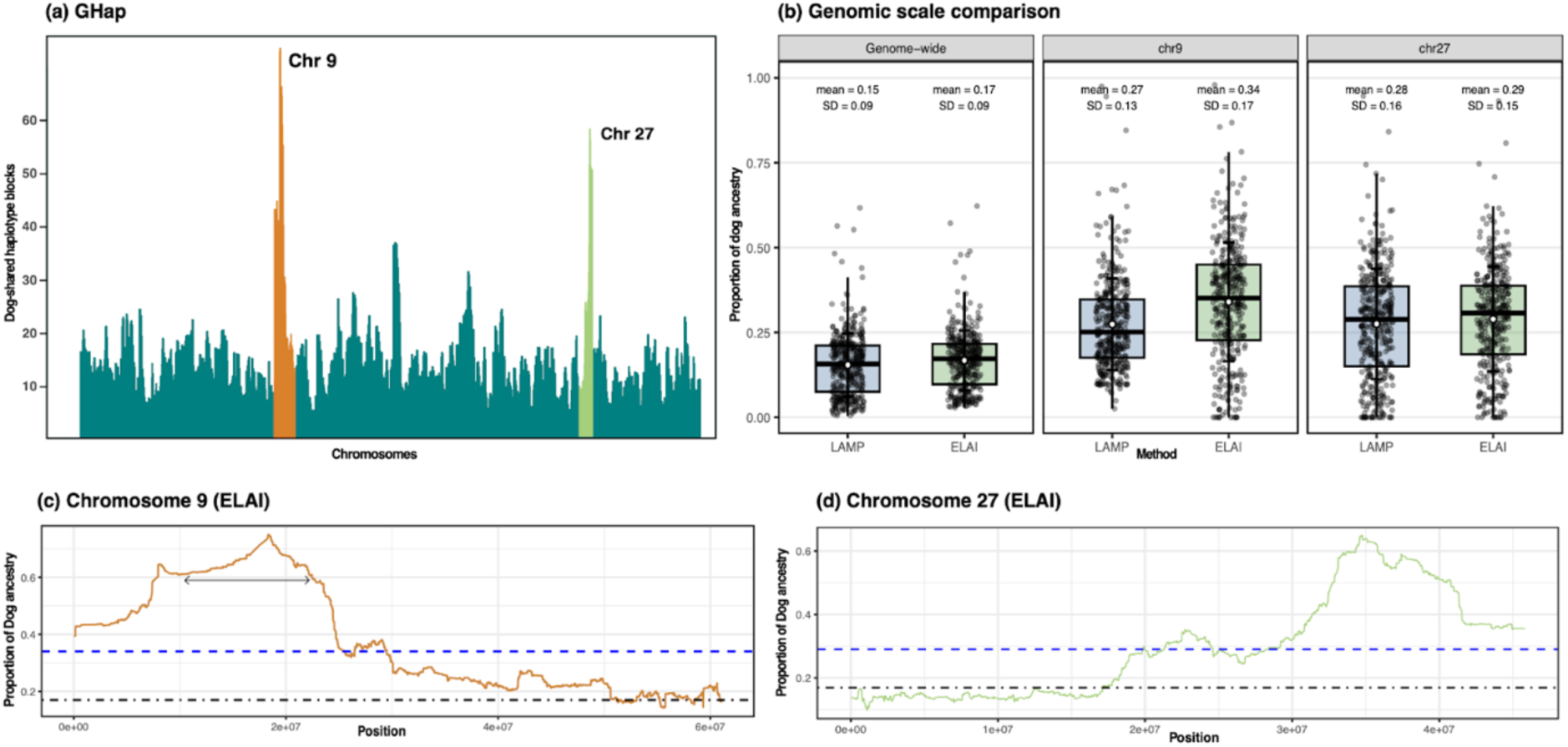
Dog ancestry across chromosomes inferred using haplotype sharing and local ancestry analyses. **(a)** Genome-wide distribution of haplotype blocks shared between dingoes and domestic dogs inferred with GHap. The y-axis indicates the number of dog-shared haplotype blocks per genomic window. Chromosomes 9 and 27 show pronounced peaks, indicating regions with elevated apparent dog ancestry. **(b)** Local ancestry inference along chromosome 9 based on ELAI. The y-axis represents the proportion of dog ancestry along the chromosome. The black dot–dash line indicates the genome-wide average introgression level, and the blue dashed line indicates the chromosome-specific mean. The region initially identified as a putative introgressed block encompasses a large chromosomal inversion previously described in domestic dogs (Field et al., 2022), whose coordinates are indicated by the arrow and likely explain the observed ancestry pattern. **(c)** Local ancestry inference along chromosome 27 inferred with ELAI. Line types and thresholds are as in panel (b). **(d)** Comparison of local ancestry estimates across genomic scales. Distribution of individual-level proportions of dog ancestry in dingoes inferred using LAMP and ELAI at three genomic scales: genome-wide, chromosome 9, and chromosome 27. Points represent individuals (jittered for visibility). Boxplots show medians and interquartile ranges; white circles indicate means and vertical bars represent ±1 standard deviation. Both mean ancestry and variance increase markedly on chromosomes 9 and 27 relative to genome-wide estimates.

#### 3.2.2. Identification of Introgressed Regions

GHap analyses confirmed these chromosomes as primary regions of dog ancestry, where distinct introgressed haplotype blocks were identified (Figure 2c-d), spanning 19.5 Mb on chromosome 9 and 8.9 Mb on chromosome 27. Notably, both chromosomes 9 and 27 exhibited elevated dog ancestry in Eastern and Southern dingoes, with chromosome 27 also showing some introgression in Central populations. These patterns suggest partial geographic structure in introgression. In addition, isolated peaks of introgression were observed on chromosomes 13, 14, and 24. These isolated peaks, though smaller in size, may represent additional regions of interest for adaptive introgression (Tables S4, S5, Figure S5).

We explored linkage disequilibrium (LD) patterns in regions of introgression on chromosomes 9 and 27, which harbored the most prominent dog-derived haplotype blocks. Although LD (R^2^) was calculated genome-wide for all chromosomes (mean = 0.103; median = 0.049; max = 1.000), we focus here on chromosomes 9 and 27 due to their relevance. On chromosome 9, the mean R^2^ was 0.069 (median = 0.020; max = 1.000), and on chromosome 27 it was 0.055 (median = 0.018; max = 0.984). Crucially, over 95% of all SNP pairs with R^2^ > 0.8 on chromosome 9 fall within our previously defined introgressed interval on that chromosome, and over 90% of high-LD SNP pairs on chromosome 27 co-localize with its candidate block. This strict spatial confinement of elevated LD to the introgressed regions indicates that recombination has been less effective at breaking down these haplotypes locally, potentially in combination with selection or recent demographic history.

To place these values in a genome-wide context, we summarized median LD (R² ≤ 500 kb) for all autosomes. Median LD values showed moderate variation across chromosomes but largely overlapped (Figure S6). Chromosome 27 fell toward the lower end of the genome-wide distribution, but did not differ significantly from the remaining autosomes when compared against chromosomal medians (Wilcoxon rank-sum test, p = 0.053).

Using a fine-scale dog recombination map (Kidd, 2026), the chromosome 27 block showed a length-weighted mean recombination rate of 1.39 cM/Mb. In a permutation test based on 10,000 length-matched windows sampled across chromosome 27, the block rate fell well within the chromosome-wide distribution (empirical p for unusually low recombination = 0.74; Figure S7), indicating that the candidate region does not coincide with a recombination coldspot. This provides an additional support for adaptive introgression as the potential cause of the elevated introgression in this chromosomal block.

The fragment on chromosome 9 initially identified as an introgressed block was later found to encompass a large chromosomal inversion previously described in domestic dogs but absent in dingoes (Field et al., 2022). Using inversion coordinates provided by the authors and converting them to the canFam6 reference genome with the UCSC liftOver tool (Raney et al., 2024), we confirmed that the inversion spans an interval fully contained within the candidate region. This region does not show decreased genetic differentiation between dingoes and domestic dogs (Figure S8), which would be expected in the case of intense introgression inferred from the local ancestry patterns (Figure 2, Figure S3a). The chromosome 9 inversion is present in most but not all dogs breeds, so for this chromosomal region dogs that do not have the inversion are more similar to dingoes than to other dogs. This shared similarity between dingoes and some dogs could have resulted in an incorrect signal of introgression in the local ancestry analysis, with dogs that do not have the inversion being interpreted as donors of this variant to dingoes (Figure S3a). A lack of the inversion in dogs is rarer than presence, which is more consistent with a scenario where dingoes are the donors of this variant to dogs. However, this in turn is inconsistent with the pattern observed in other parts of chromosome 9 (no introgression from dingoes to dogs), which could be the reason why introgression from dogs to dingoes was inferred. Field et al. (2022) showed that a lack of the inversion is an ancestral state (occurring in grey wolves and dingoes) and the inversion occurred recently during the dog breed diversification process. Therefore, it is unlikely that the pattern observed in chromosome 9 reflects actual gene flow from dogs to dingoes, and we excluded this chromosomal region from subsequent introgression analyses.

Besides the regions with elevated dog ancestry, we also detected dog ancestry deserts on eight chromosomes (Table S4, Figure S9). Most of these ancestry deserts included protein-coding genes, indicating possible targets of negative selection. Notably, one of these regions included Bone Morphogenetic Protein 4 (BMP4), a gene with well-established roles in developmental processes (Ye et al., 2022). These ancestry deserts were generally short, suggesting that strong, genome-wide barriers to introgression are limited. Dog-enriched variants within ancestry deserts were predominantly non-coding (upstream, downstream, or intronic) and showed no evidence of deleterious protein-coding changes. Across 21 variants under our primary filter and 25 variants under a more permissive filter, we detected no missense or predicted loss-of-function variants.

#### 3.2.3. Functional Analysis of Genes Within Introgressed Regions

Within the introgressed block on chromosome 27, most genes showed signatures of purifying selection based on dN/dS ratios significantly lower than 1 (Figure 4; Table S5). One gene, OR8S3 (Olfactory receptor family 8 subfamily S member 8), exhibited a dN/dS ratio significantly above 1, consistent with positive selection (p = 0.0012). Additional introgressed regions on chromosomes 13, 14, and 24 also contained genes under purifying selection. Specifically, GNPDA2 and KCTD8 (chr13), and SDHAF3 (chr14) showed significant results, while ASNS, EMILIN3, and CHD6 were non-significant or marginally so. These analyses were based exclusively on coding sequences; therefore, potential selective pressures on non-coding or regulatory variants within the introgressed regions cannot be assessed with this approach. Although the chromosome 9 region was later shown to represent an inversion rather than an introgressed block, we nonetheless examined its protein-coding genes and found that two of them -GJC1 (Gap Junction Protein Gamma 1) and TCAP (Titin-Cap)-exhibit signals of positive selection (Table S6).

#### 3.2.4. Phylogenetic and Genome-Wide Evidence of Introgression

For clarity of visualization and interpretation, the main phylogenies shown in Figure 3 were reconstructed using a representative subset of five randomly selected individuals per population or species, except for NGSD (n = 2) and the golden jackal (n = 1). The phylogenies reconstructed using all available individuals are provided in the Supplementary Material (Figure S10). The Maximum Likelihood tree reconstructed with IQ-TREE based on genome-wide SNP data revealed reciprocally monophyletic clades for dingoes and domestic dogs (with domestic dog cluster grouping purebred and mixed-breed dogs independent of geographic origin as well as European free-ranging dogs). In contrast, New Guinea Singing Dogs (NGSD) did not form a reciprocally monophyletic clade distinct from dingoes, instead clustering closely with them-consistent with their shared evolutionary history (Figure 3a, Figure S10a). The phylogenetic tree for the introgressed region on chromosome 27 revealed clustering between domestic dogs and dingoes, consistent with introgression (Figure 3b, Figure S10b). All populations except Big Desert and captive dingoes had introgressed haplotypes in their gene pool, and these haplotypes clustered with different European dog clades, implying that the introgression is a result of multiple dingo-dog cross-breeding events. Domestic dogs did not have any haplotypes clustering with dingo-specific haplotypes, implying no introgression from dingoes to dogs in this chromosomal region.

**Figure 3.**
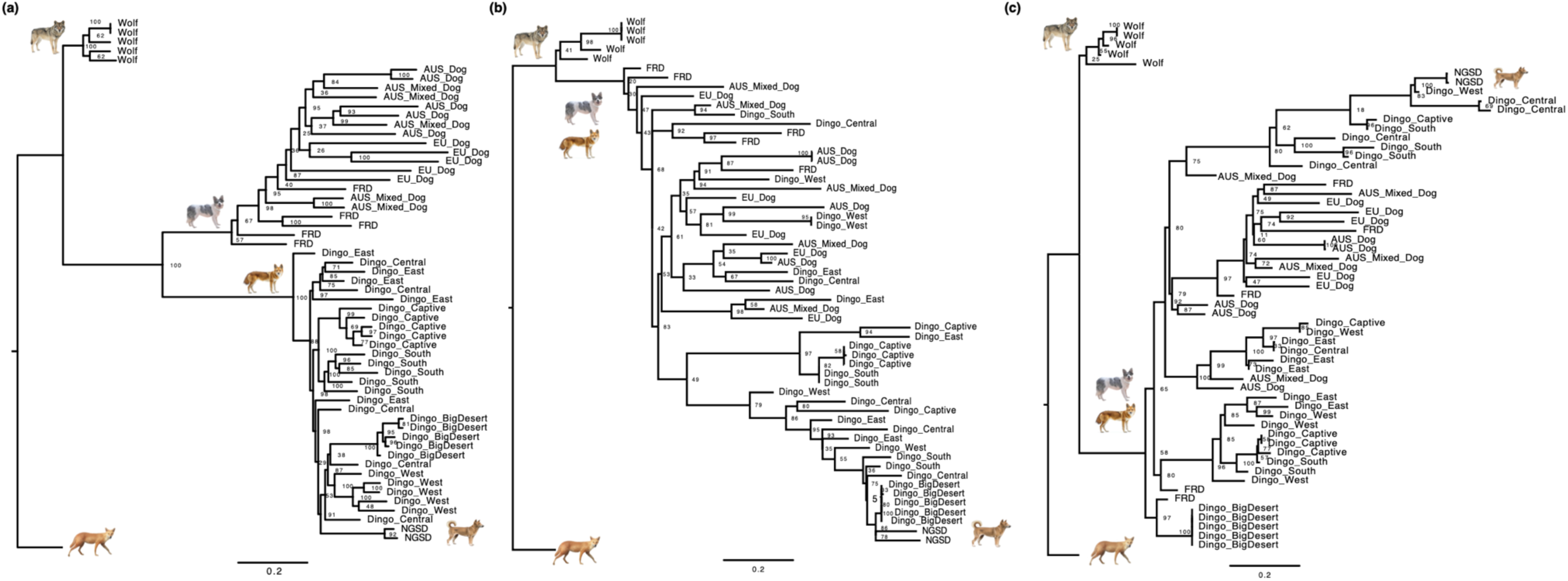
Phylogenetic analyses. **(a)** Maximum Likelihood phylogeny reconstructed with IQ-TREE using genome-wide SNP data from a subset of five randomly selected individuals per population or species, except for New Guinea Singing Dogs (NGSD; n = 2) and the golden jackal (outgroup; n = 1). The tree shows species- and population-level clustering. Ultrafast bootstrap support values (UFBS) are shown at the nodes, and the scale bar represents the expected number of nucleotide substitutions per site. **(b)** Maximum Likelihood phylogeny for the introgressed block on chromosome 27, using the same subset of individuals. Clustering of domestic dogs and dingoes in this region is consistent with localized introgression. The full phylogenies including all individuals are provided in the Supplementary Material (Figure S10 a-c). **(c)** Chromosome 9 inversion region: ML phylogeny built on SNPs within the inversion detected on chromosome 9. Some European domestic dogs cluster closely with dingoes mirroring the pattern from LAMP-LD local ancestry analyses.

**Figure 4.**
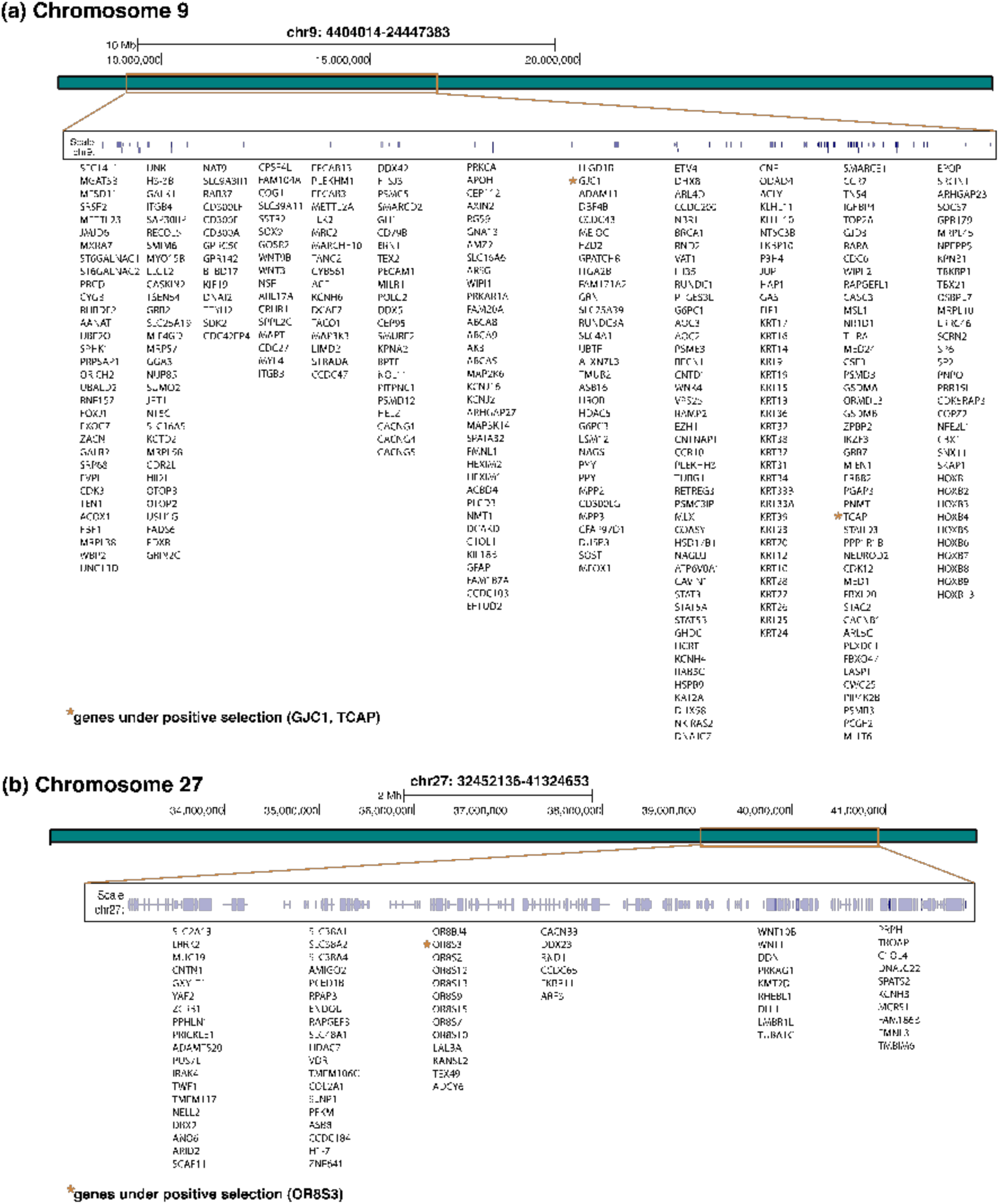
Genomic regions of interest in Chromosomes 9 and 27. **(a)** Detailed view of Chromosome 9, highlighting genes located within the inverted region that initially appeared as an introgressed block. Two genes-GJC1 and TCAP-showed signals of positive selection and are marked with an asterisk (*). **(b)** Detailed view of Chromosome 27, where a well-defined introgressed block was identified. One gene-OR8S3-was under positive selection, also indicated by an asterisk (*). Most genes in both regions were under purifying selection, with dN/dS ratios below 1. For a full list of genes, selection statistics, and associated p-values, see Table S5.

In the phylogeny for the chromosome 9 region, some European domestic dogs clustered closely with dingoes (Figure 3c, Figure S10c), while other dogs formed distinct clusters. This pattern is consistent with the presence of a chromosomal inversion previously described in some, but not all, dog breeds and absent in dingoes (Field et al. 2022). As such, the phylogenetic signal in this region likely reflects haplotype similarity maintained by suppressed recombination within the inversion, rather than introgression.

Additionally, ABBA-BABA analyses (D-statistics) provided strong genome-wide evidence of introgression from domestic dogs into dingoes. Across all dingo populations, the D-statistic was 0.1547 (Z = 7.20, p < 1×10⁻¹²), with an f₄-ratio of 0.192, indicating a substantial excess of shared derived alleles between dingoes and domestic dogs (Table 2). When examined individually, all dingo populations exhibited positive and highly significant D-statistics, with the strongest signal detected in Eastern populations (D = 0.193, Z = 8.53, p = 2.3×10⁻¹⁶), followed by those from the South (D = 0.179, Z = 7.08, p = 1.5×10⁻¹²), congruent with our local ancestry results (Table 2). These findings underscore geographic variation in the intensity of dog ancestry across the dingo range. Focusing on the introgressed region on chromosome 27, the signal of dog ancestry was even stronger (D = 0.263, Z = 3.35, p = 0.0008), with an f₄-ratio of 0.556, suggesting substantial localized introgression. Among populations, only trios with informative ABBA-BABA sites and p < 0.05 were reported. Central dingoes exhibited the strongest signal (D = 0.282, Z = 3.58, p = 0.0003), followed by those from the South and West (D = 0.235 and 0.210; Z = 2.83 and 2.69; p = 0.0046 and 0.0072, respectively). Across the chromosome 27 candidate block, f_d values were consistently elevated across most genomic windows (typically ranging from ∼0.4 to >0.6), indicating a strong excess of dog-derived ancestry concentrated within this region. In contrast, f_d values decreased sharply outside the core of the block, supporting the interpretation that introgression on chromosome 27 is spatially localized rather than driven by genome-wide background processes.

**Table 2.**
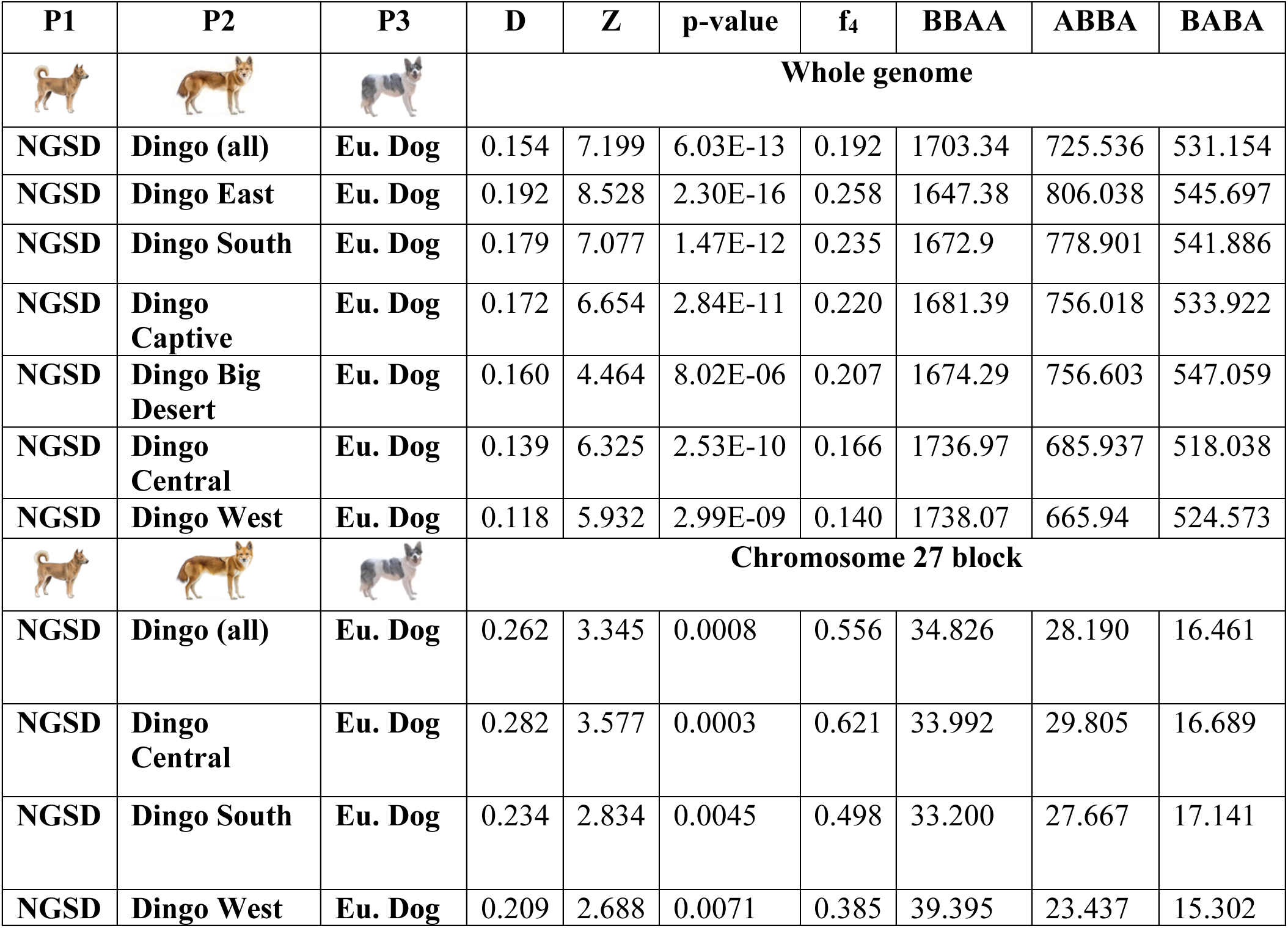
Summary of ABBA–BABA (D-statistic) tests of introgression between dingoes and European dogs, conducted with Dsuite on (i) whole-genome VCF data and (ii) a candidate introgressed block on chromosome 27. In each test, P1 was the New Guinea singing dog (NGSD; the sister lineage to dingoes), P2 was either all dingoes combined or one of six regional dingo populations (East, South, Captive, Big Desert, Central, West), and P3 was the pure breed European dog samples (Eu. Dog), with the jackal as an outgroup. For each trio we report Patterson’s D (excess allele sharing between P2 and P3 versus P1 and P3), its Z-score (block-jackknife), two-tailed p-value, the f₄-ratio (proportion of P3 ancestry in P2), and counts of BBAA (derived allele shared by P1+P2), ABBA (shared by P2+P3) and BABA (shared by P1+P3) site patterns. In the genome-wide analyses, all D values are significantly positive (p < 0.001), indicating gene flow between dingoes and European dogs. For chromosome 27, only trios with informative ABBA/BABA sites and p < 0.05 are shown. Elevated D and f₄-ratio values on chromosome 27 relative to genome-wide averages highlight a localized introgression signal in this candidate region.

| P1 | P2 | P3 | D | Z | p-value | f<sub>4</sub> | BBAA | ABBA | BABA |
| --- | --- | --- | --- | --- | --- | --- | --- | --- | --- |
|  |  |  | Whole genome |  |  |  |  |  |  |
| NGSD | Dingo (all) | Eu. Dog | 0.154 | 7.199 | 6.03E-13 | 0.192 | 1703.34 | 725.536 | 531.154 |
| NGSD | Dingo East | Eu. Dog | 0.192 | 8.528 | 2.30E-16 | 0.258 | 1647.38 | 806.038 | 545.697 |
| NGSD | Dingo South | Eu. Dog | 0.179 | 7.077 | 1.47E-12 | 0.235 | 1672.9 | 778.901 | 541.886 |
| NGSD | Dingo Captive | Eu. Dog | 0.172 | 6.654 | 2.84E-11 | 0.220 | 1681.39 | 756.018 | 533.922 |
| NGSD | Dingo Big Desert | Eu. Dog | 0.160 | 4.464 | 8.02E-06 | 0.207 | 1674.29 | 756.603 | 547.059 |
| NGSD | Dingo Central | Eu. Dog | 0.139 | 6.325 | 2.53E-10 | 0.166 | 1736.97 | 685.937 | 518.038 |
| NGSD | Dingo West | Eu. Dog | 0.118 | 5.932 | 2.99E-09 | 0.140 | 1738.07 | 665.94 | 524.573 |
|  |  |  | Chromosome 27 block |  |  |  |  |  |  |
| NGSD | Dingo (all) | Eu. Dog | 0.262 | 3.345 | 0.0008 | 0.556 | 34.826 | 28.190 | 16.461 |
| NGSD | Dingo Central | Eu. Dog | 0.282 | 3.577 | 0.0003 | 0.621 | 33.992 | 29.805 | 16.689 |
| NGSD | Dingo South | Eu. Dog | 0.234 | 2.834 | 0.0045 | 0.498 | 33.200 | 27.667 | 17.141 |
| NGSD | Dingo West | Eu. Dog | 0.209 | 2.688 | 0.0071 | 0.385 | 39.395 | 23.437 | 15.302 |

#### 3.2.5 Population-level variation in introgression revealed by Bayesian genomic clines

Bayesian genomic cline analyses revealed pronounced population-level differences in the introgression patterns among dingo populations. Estimates of individual hybrid indices (HI) confirmed substantial heterogeneity in overall levels of dog ancestry, with eastern and southern populations showing higher mean HI values, whereas Western and Big Desert populations exhibited lower average introgression (Table S7).

The populations also varied markedly in locus-specific introgression patterns. Eastern and southern dingoes showed relatively low dispersion of genomic cline centers (SDc) and slopes (SDv), indicating more homogeneous ancestry transitions across loci. In contrast, Western and Central populations exhibited higher SDc values, consistent with greater locus-specific heterogeneity in introgression across the genome. Captive dingoes showed intermediate levels of dispersion, reflecting mixed ancestry patterns. Together, these results indicate that introgression in dingoes is not only geographically structured in magnitude but also differs in its genomic architecture among populations. Overall, populations with higher mean HI showed comparatively lower SDc/SDv, whereas populations with lower mean HI showed higher dispersion, although this pattern was not uniform across all populations. These patterns are consistent with results from local ancestry inference and genome-wide introgression analyses, reinforcing the conclusion that introgression dynamics vary spatially across Australia.

### 3.3 Influence of Environmental Variation on Spatial Introgression

Mantel tests revealed no significant relationship between genetic and geographic distances (Figure S11a, r = 0.027, p = 0.1913). However, a positive correlation between genetic and environmental distances (Figure S11b, r = 0.1515, p < 0.0001) supports genetic differentiation driven by environmental gradients.

To identify the environmental variables most relevant to introgression patterns and reduce redundancy among predictors, we conducted a partial Redundancy Analysis (pRDA) as an intermediate step. This analysis revealed that environmental factors significantly contribute to genetic differentiation between dingo populations, with the full model (climate, geography, and population structure) explaining 9.7% of the genetic variation (p < 0.001; Table 3). Climate-specific variables were the strongest predictors, accounting for 4.9% of the variation, while population structure and geography contributed 3.0% and 1.8%, respectively. Among the environmental predictors, the maximum temperature of the warmest month (BIO5), mean temperature of the driest quarter (BIO9), and the Human Footprint Index (HFPI) emerged as the most influential variables (Figure 5).

**Figure 5.**
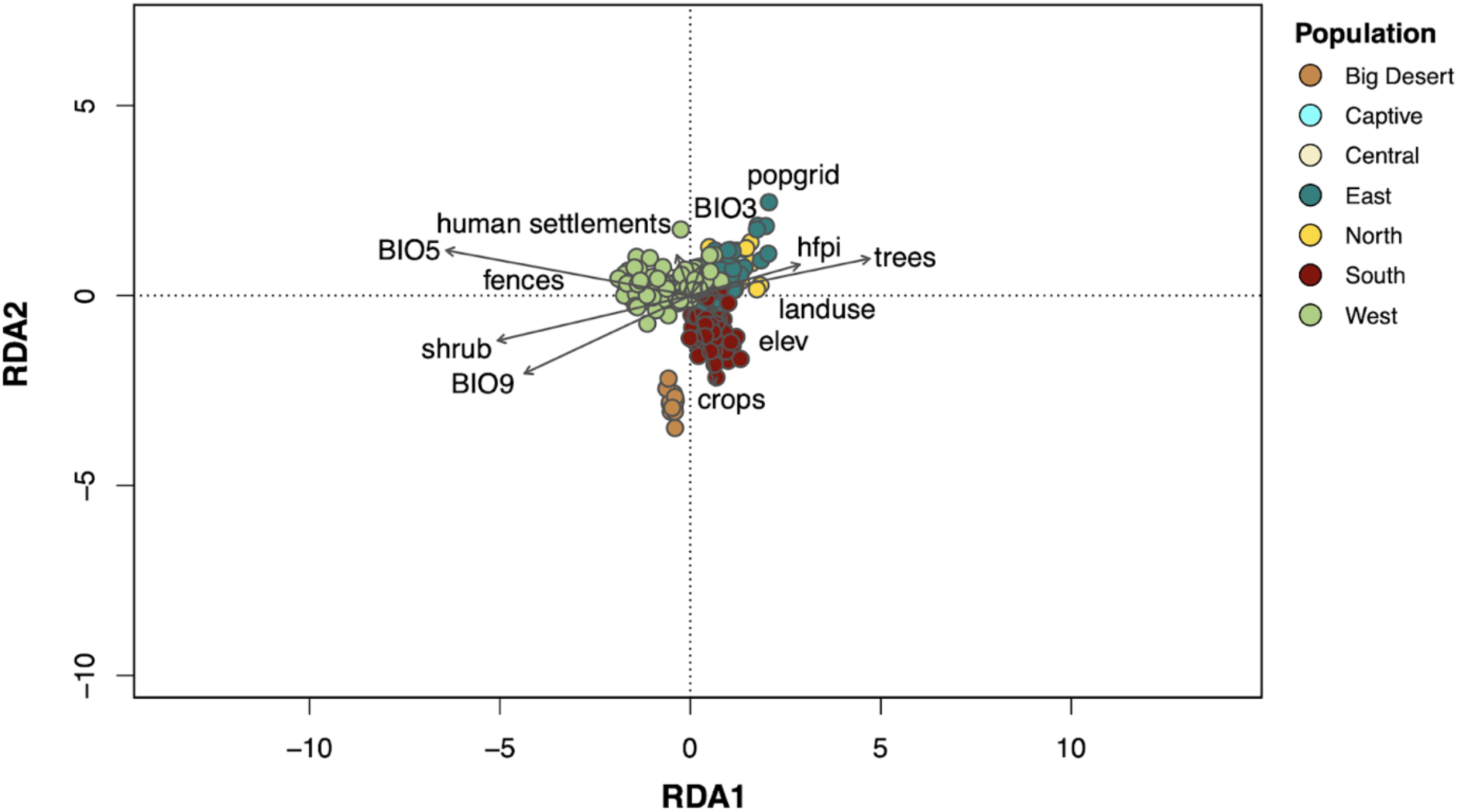
Environmental drivers of genetic variation. Redundancy Analysis (RDA). Each point is an individual colored by population (Big Desert, Central, East, North, South, West, Captive). Environmental predictors are shown as arrows: popgrid: regional human population density, human_settlements: proximity to built-up areas, trees: forest cover, fences: livestock fencing density, landuse: proportion of modified land types, crops: agricultural land cover, shrubs: shrubland cover, elev: elevation, BIO3, BIO5, BIO9: bioclimatic variables (isothermality, max temperature of warmest month, mean temperature of driest quarter, respectively). Arrow direction and length indicate the variables’ loadings on RDA1 and RDA2; longer arrows denote stronger associations with dingo genetic structure.

**Table 3.**
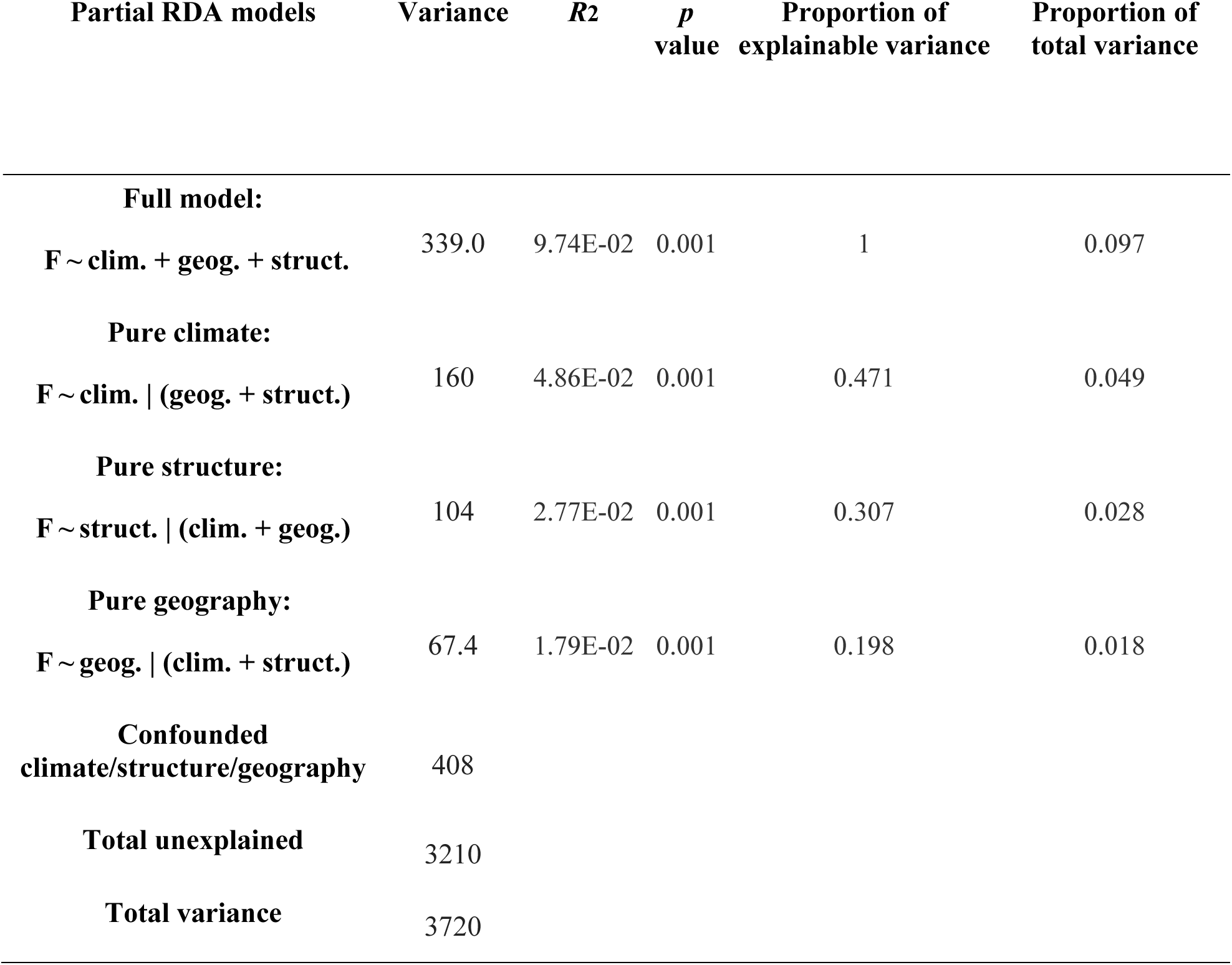
Results of partial Redundancy Analysis (partial RDA) examining the influence of climate, genetic structure, and geography on genetic variation. The analysis decomposes the variation into components attributable to climate, genetic structure, and geography. The proportion of explainable variance represents the total constrained variation explained by the full model.

In addition to identifying key environmental predictors, the pRDA highlighted loci potentially involved in local adaptation. Across the second and third constrained axes, 524 outlier loci were associated with environmental variables, particularly temperature-related factors (e.g., BIO5: 414 loci; BIO9: 353 loci) and vegetation-related features such as shrub and tree cover (388 and 386 loci, respectively; Table S2e). These loci suggest possible genetic responses to climatic and landscape features, supporting the hypothesis of environmental selection pressures shaping genetic differentiation.

To directly evaluate how environmental variation relates to the proportion of dog introgression, we used Random Forest models with individual-level dog ancestry as the response variable. Across all individuals, introgression proportions ranged from 0.03 to 0.48, with a mean of 0.16, and with substantial heterogeneity among populations. The global Random Forest model explained a substantial proportion of variance in introgression (out-of-bag R² = 0.66; MSE = 0.0022). Variable importance rankings consistently identified climatic predictors as the strongest contributors, particularly maximum temperature of the warmest month (BIO5), precipitation of the coldest quarter (BIO19), and mean temperature of the wettest quarter (BIO8), followed by additional temperature and precipitation-related variables (Figure S12). Anthropogenic predictors, including HFPI and land-use metrics, contributed more modestly but consistently to model performance.

Predictive performance varied markedly among populations when evaluated using a leave-one-population-out cross-validation approach. Model performance was highest in southern populations (R² = 0.69) and moderate in Western and Northern populations (R² = 0.34 and 0.32, respectively), whereas predictive power was very low in Eastern, Central, and Big Desert populations (R² ≤ 0.03). This spatial heterogeneity indicates that relationships between environmental conditions and introgression are not uniformly transferable across regions.

To place these results in a biological context, we quantified effect sizes along observed environmental gradients. Shifting BIO5 from the 10th to the 90th percentile of its empirical range was associated with an average decrease of approximately 4.7% in predicted dog ancestry, whereas an equivalent shift in BIO19 corresponded to an increase of approximately 3.9%. Changes along BIO8 produced smaller effects (≈1%). These results indicate that individual climatic gradients exert moderate effects on introgression levels, but their combined influence can generate substantial spatial variation in hybridization across dingo populations.

## 4. Discussion

### 4.1 Introgression Patterns in Australian Dingoes

Our study provides new insights into the genomic patterns of introgression between dingoes and domestic dogs in Australia, emphasizing the complex interplay between gene flow and environmental adaptation. By integrating high-resolution genomic data with ecological variables, we highlight how environmental and anthropogenic factors are influencing the spatial and genomic distribution of dog introgression in dingo populations. Population structure analyses, including PCA and ADMIXTURE, confirmed regional differentiation among dingo populations reported in prior studies (Cairns et al., 2023, Souilmi et al., 2024) as well as strong genetic differentiation between dingoes and domestic dogs, with minimal overlap between clusters. Nonetheless, admixture signals were detected, particularly in dingoes from Eastern and Southern Australia. Local ancestry inference methods (LAMP-LD, ELAI, and GHap) further revealed complex and heterogeneous introgression signatures across the genome. ABBA–BABA analyses also supported the presence of genome-wide introgression, albeit with varying intensity across geographic regions. Bayesian genomic cline analyses further indicate that populations differ not only in the extent of introgression but also in its genomic architecture, with some populations showing homogeneous ancestry transitions across loci and others exhibiting pronounced locus-specific heterogeneity. Genomic regions with overrepresented dog ancestry were identified on four chromosomes, most prominently on chromosome 27. Importantly, this localized signal was also supported by elevated f_d values across the chromosome 27 block, a statistic specifically designed to detect introgression at fine genomic scales and to reduce the confounding effects of incomplete lineage sorting (Martin et al., 2015). Taken together, our findings suggest historical and ongoing dog introgression, with hybridization cases occurring infrequently but leading to backcrossing, which has left variable genomic footprints across the dingo genome.

Differences in estimated ancestry proportions across studies likely reflect contrasts between global clustering approaches and locus-specific ancestry inference methods. Our global ancestry estimates are broadly consistent with those reported by Cairns et al. (2023), whereas higher admixture proportions emerge when applying local ancestry inference. In contrast, global clustering methods applied by Weeks et al. (2024) revealed minimal or no admixture. All the methods we applied were consistent in showing the presence of dog admixture in dingoes, although its proportions differed between the global ancestry estimate using ADMIXTURE (9% at average) and local ancestry estimates (12-15%). All the methods were also consistent in identifying the lowest admixture proportions in the Central and Western populations and the highest proportions in the Eastern, Southern and Captive populations. Our local ancestry estimates are in line with estimates based on whole genome sequences using an ancient DNA baseline (Scarsbrook et al., 2025). Very low (<0.0015) dingo admixture proportions in European dogs inferred based on local ancestry methods show that the false positive rate is low. In contrast, ADMIXTURE identified 3% of dingo ancestry in European free-ranging dogs, which can be attributed to recent shared ancestry. These differences highlight how global clustering approaches may dilute or underestimate localized introgression signals, whereas local ancestry inference has greater power to detect both contemporary and historical gene flow. However, when historical introgression has become pervasive across the genome, it may be interpreted as ancestral variation rather than discrete admixture tracts. Local ancestry methods may be thus considered as more precise, yet they may be prone to other sources of errors, such as those resulting from the structural changes occurring in the process of dog evolution (see below).

Our integrative approach, combining local ancestry methods with phylogenomic analyses, enabled us to detect introgression signals that were not apparent in previous SNP-based studies. A key strength of our work lies in its broad geographic sampling, allowing for a comprehensive assessment of hybridization dynamics across the continent. Below, we discuss the environmental and anthropogenic factors shaping introgression patterns, highlight genomic regions of interest that may reflect adaptive introgression, and consider the implications of these findings for dingo conservation and genetic monitoring.

### 4.2 Environmental and Anthropogenic Drivers of Hybridization

Human-modified landscapes likely play a key role in facilitating hybridization between Australian dingoes and domestic dogs. Indeed, our results detected significant interactions between proximity to human settlements and introgression, which likely arise from higher densities of free-roaming owned dogs (Sparkes et al., 2022), more intense lethal control measures, or the effects of landscape alteration. Local ancestry analyses using LAMP-LD and ELAI reveal clear geographic patterns of introgression, with dingoes in the Central, Western, and Big Desert regions exhibiting considerably lower levels of dog ancestry compared to those in the East and South. Notably, some of the differentiation between Western and Eastern dingo populations predates European colonization, indicating long-standing historical isolation alongside recent admixture (Souilmi et al., 2024). These findings are consistent with population structure results and support the idea that introgression varies geographically across dingo populations. While population structure likely reflects multiple historical and demographic processes, differences in introgression patterns appear to contribute to the observed genetic clustering. This interpretation aligns with previous genomic studies (Stephens et al., 2022; Cairns et al., 2023; Weeks et al., 2024) and suggests that ecologically and geographically isolated populations-such as those in the Central region-experience minimal admixture, likely due to both reduced contact with domestic dogs and harsher environmental conditions that limit their overlap. In contrast, dingoes from Eastern and Southern Australia – regions with higher human footprint - show higher levels of introgression.

Notably, our results revealed a strong signal of isolation by environment (IBE) but no significant evidence of isolation by distance (IBD) in the present dataset. Although previous studies have reported IBD patterns in dingoes (Stephens et al., 2022; Weeks et al., 2024), our findings suggest that environmental heterogeneity may play a particularly important role in shaping both population divergence and introgression at the spatial and genomic scale examined here. In other canids, such as wolves, long-distance dispersal is common, but environmental gradients remain key determinants of gene flow (Geffen et al., 2004; Pilot et al., 2006; Leonard et al., 2014). In dingoes, this interplay between dispersal potential and environmental variation may similarly create localized barriers to gene flow, which influence both adaptation and introgression outcomes.

Understanding the environmental and anthropogenic drivers of introgression is essential to clarifying how hybridization shapes dingo populations in increasingly altered landscapes. Dingoes inhabit a wide range of environments, and while direct assessments remain limited, interactions between human activity and climatic conditions likely create varying opportunities for contact and genetic exchange with domestic dogs (Cairns et al., 2020). For example, ecological overlap in peri-urban areas and pastoral lands may increase dingo-dog encounters, particularly where human-associated resources concentrate domestic dogs in areas that also attract dingoes. These landscapes not only promote higher contact rates but also impose selective pressures that may favor the retention of dog-derived alleles adaptive in human-dominated landscapes in dingo genomes.

Our Random Forest (RF) analyses highlight the importance of environmental gradients, together with anthropogenic factors, in driving these patterns. Climatic variables emerged as the strongest predictors of introgression, particularly maximum temperature of the warmest month (BIO5) and precipitation-related variables (e.g. BIO19), suggesting that climate plays a central role in shaping introgression dynamics. The Human Footprint Index (HFPI) also contributed consistently to model performance, supporting the idea that hybridization and subsequent introgression may be facilitated in areas with greater human disturbance. Together, these results indicate that introgression patterns are strongly structured by climate, with human influence acting as a context-dependent modifier. Additionally, climatic variables such as maximum temperature of the warmest month (BIO5) and other temperature- and precipitation-related factors likely shape hybridization dynamics by influencing habitat suitability, movement patterns, and resource use. In Australia, climatic conditions strongly shape patterns of human settlement, with milder climates supporting higher human densities. Because domestic dog density closely tracks human density (Gompper, 2014), climate may indirectly modulate dingo-dog encounter rates by structuring dog abundance and human land use. In arid or seasonally dry regions, dingoes may concentrate around permanent water sources, pastoral infrastructure or areas with anthropogenic food subsidies, potentially increasing contact with dogs during climatically stressful periods.

Dependence of hybridization rates on environmental variables have been reported across a wide range of taxa, indicating that environmentally mediated hybridization is a general phenomenon rather than a system-specific exception. In birds, human habitat disturbance has been shown to increase hybridization rates between closely related species, as observed in black-capped and mountain chickadees, where landscape modification alters contact zones and mating opportunities (Grabenstein et al., 2023). In freshwater fishes, hybridization outcomes among trout species vary predictably with environmental context and historical demographic processes linked to habitat alteration and management history (Mandeville et al., 2019). Similarly, asymmetrical hybridization and spatial genetic structure in killifish hybrid zones have been shown to reflect the combined influence of environmental gradients and landscape features (Hardy et al., 2021). In plants, environmental heterogeneity and landscape structure can shape mosaic hybrid zones and influence the maintenance of reproductive barriers (Faske et al., 2024). Together, these studies reinforce the view that hybridization rates and genomic outcomes are strongly contingent on extrinsic environmental conditions, paralleling the patterns we observe in dingoes. Within canids, similar environmentally mediated patterns of admixture have also been reported. Studies in North America have demonstrated that urbanization and landscape changes significantly influence admixture between coyotes and wolves (Stronen et al., 2012), underscoring the importance of human-mediated pressures in shaping hybridization outcomes in wild canids (Pilot et al., 2021).

### 4.3 Adaptive Significance of Introgressed Haplotype Blocks

Building on the geographic patterns of introgression revealed through our analyses, we identified a major introgressed block on chromosome 27 and several smaller peaks on chromosomes 13, 14, and 24. These regions may represent portions of the dingo genome where dog-derived alleles have been retained by selection acting on functionally relevant genes. In most genes within the chromosome 27 block and in the other smaller peaks, dN/dS values are below 1, suggesting purifying selection. We note, however, that historical bottlenecks and reduced effective population size can relax purifying selection and alter expectations for dN/dS ratios (Kryazhimskiy and Plotkin, 2008), which cautions against interpreting elevated dN/dS values as definitive evidence of selection. Historical bottlenecks experienced by dingoes (Kumar et al., 2023) can generate heterogeneous genomic patterns and accelerate the fixation of haplotypes, particularly in regions of low recombination. In addition, heterogeneity in recombination rates across chromosomes is known to shape genomic landscapes independently of selection (Burri et al., 2015), and neutral variation in introgression rates has been widely documented in genomic cline analyses (Gompert and Buerkle, 2011).

Although the signatures of natural selection should be treated with caution before they can be tested at the phenotypic level, we detected one potential case of adaptive introgression: an olfactory receptor gene, OR8S3, within the chromosome 27 block, exhibits a clear signature of positive selection, suggesting that the introgressed allele may confer a sensory advantage. Olfactory receptors (ORs) are central to foraging, social communication, and environmental sensing in mammals (Robin et al., 2009), and wild canids often show strong selection on OR repertoires in response to ecological pressures. Comparative data indicate that dingoes maintain a larger, wolf-like OR repertoire than modern dog breeds-likely reflecting continued reliance on prey hunting (Mouton et al., 2025). In this context, the retention and selection of a dog-derived OR8S3 variant could represent a fine-tuned sensory adaptation, perhaps aiding dingoes in detecting novel anthropogenic cues or exploiting human-associated resources.

A region on chromosome 9 was flagged by all local-ancestry methods, but exhibited an unusual introgression pattern (Figure S3a). We found this signal arose from a large inversion segregating in dogs (Field et al., 2022) that suppresses recombination and conserves extended haplotypes. Because this inversion is neither fixed in dogs nor present in dingoes (Field et al., 2022), we interpret the shared haplotypes as retained ancestral variation rather than true gene flow and therefore we have excluded this region from our introgression analyses.

Linkage disequilibrium (LD) analyses of chromosomes 9 and 27 further revealed heterogeneous patterns. Haplotype blocks identified as introgressed from dogs through local ancestry inference coincided with regions of strong LD. This elevated LD may be due to reduced recombination rates, which can arise from structural features such as inversions (as seen on chromosome 9) or from selective sweeps maintaining favorable haplotypes. In contrast, LD was generally lower outside the introgressed blocks, consistent with background recombination eroding ancestral haplotypes over time.

In addition to introgressed haplotype blocks, we also identified eight ancestry deserts across the genome (Table S6), defined as regions consistently reduced of dog ancestry. The limited number and generally small size of these deserts indicate that resistance to dog introgression does occur in the dingo genome but is relatively uncommon. This pattern is consistent with the recent divergence between dingoes and domestic dogs, which likely constrains the accumulation of strongly deleterious introgressed variants. Supporting this interpretation, variants located within ancestry deserts were predominantly non-coding, and we detected no missense or predicted loss-of-function mutations within these regions. The functional and evolutionary significance of these ancestry deserts, including the roles of individual genes located within them, remains an important topic for future investigation. Together, these results suggest that purifying selection against dog-derived alleles is present but localized, acting on specific genomic regions rather than imposing broad barriers to introgression.

### 4.4 Comparative Insights from Other Canids

Comparative evidence from other canids highlights the potential for adaptive introgression to enhance resilience in challenging environments. In North America, for example, introgression has enabled local adaptation in coyotes, where wolf-derived alleles contribute to ecological versatility (vonHoldt et al., 2016). In Tibetan dogs, introgression of EPAS1 alleles from high-altitude wolves facilitates hypoxia tolerance (Miao et al., 2017; Wang et al., 2014), while European wolves may exhibit increased tolerance to human-modified landscapes following hybridization with domestic dogs (Pilot et al., 2021; Sarabia et al. 2025; Lobo et al. 2025). Although the functional consequences remain to be fully quantified, the dog-derived alleles identified in dingoes may similarly confer adaptive benefits that improve survival in fragmented or anthropogenically modified habitats. At the same time, similar as in our study, studies in other species have identified genomic regions resistant to introgression (ancestry deserts), often associated with purifying selection against foreign alleles (Kim et al., 2018).

In a broader evolutionary perspective, our results emphasize that introgression can simultaneously represent a source of beneficial genetic diversity and a threat to genomic integrity. We also provide strong evidence that anthropogenic habitat modifications (quantified as Human Footprint Index) are one of key factors enhancing introgression from domestic into wild animals, which may both facilitate their adaptation to human-modified landscapes and compromise their function in natural ecosystems. By deepening our understanding of these processes, we can better assess how introgression contributes to adaptation in the face of rapid environmental change and growing anthropogenic pressures.

### 4.5 Conservation Implications and Management Strategies

The intricate relationship between introgression and environmental adaptation presents both critical challenges and unique opportunities for conserving Australian dingo populations. While introgressed genes under positive selection may confer adaptive benefits-such as enhanced survival in human-dominated landscapes-these potential advantages must be carefully weighed against their ecological consequences. Notably, the acquisition of traits that facilitate utilization of human-dominated areas and consumption of anthropogenic food could alter the dingo’s functional role as an apex predator, potentially disrupting ecosystem dynamics. In our dataset, signals of adaptive introgression appear limited and unlikely to compromise the functional distinctiveness of dingoes. Moreover, we detected eight ancestry deserts, suggesting that deleterious dog-derived alleles are likely purged by purifying selection but occur infrequently and are rapidly eliminated in early-generation hybrids. Large population sizes are required for efficient natural selection, therefore attempts to limit dog introgression by lethal control may instead intensify the problem.

From a conservation perspective, the Central population exhibited the lowest admixture across all methods, underscoring its value for conservation and as a reference “pure” dingo in comparative studies. Conservation strategies must therefore adopt spatially explicit frameworks that prioritize maintaining genetic diversity and the genetic integrity of dingo lineages while acknowledging that introgression may not pose a major threat. Importantly, decisions regarding population management-such as lethal control-should be guided by a nuanced understanding of genetic ancestry and the functional relevance of introgressed traits, rather than simplistic purity thresholds. Ultimately, by striking this balance, we can support the long-term viability of dingoes across Australia’s diverse ecological landscapes and inform evidence-based approaches to mitigate the genetic and ecological impacts of hybridization in wild canids.

## Supporting Information

Supporting information can be found online in the Supporting Information section. It includes the following files: a PDF file containing Supplementary Figures S2-S8, S11, S12, and Supplementary Table 7 as well as all Supplementary Figure and Table legends; Supplementary Tables S1-S6 as separate Excel files, and Supplementary Figures S1, S3, S9 and S10 as separate PDF files.

## Author contributions

MP and TN designed research, KMC performed lab work, BJN, KC and TMN contributed the samples, AFM collected environmental data, COM and KD analyzed data, COM wrote the manuscript, all the authors contributed to the manuscript revision.

## Data Accessibility

The dataset that support the findings of this study, including PLINK files, as well as an HTML file containing reproducible scripts and code used for the analyses, are openly available at FigShare: https://figshare.com/s/0ef4bf5568234b8ad40f. The repository contains a README file to guide users through the dataset structure and content, ensuring the analyses can be reproduced accurately.

## Acknowledgements

We thank Prof. Josephine Pemberton and two anonymous reviewers for their constructive comments on the manuscript.

