## Supplementary Material for "Domestic dog introgression in Australian dingoes: environmental drivers and evolutionary consequences"

**Title:**

**Running Title:** Causes and consequences of dog introgression in dingoes

### Supplementary Tables legends

**Table S1.** Dog and dingo data samples included in this study, providing geographic and population metadata. Columns detail each sample's population designation, breed, specific location (state, latitude, longitude), and placeholder fields (NA) where information was unavailable (attached as xls).

**Table S2.** Comprehensive environmental data and additional details (attached as xls): (a) Complete environmental dataset used in the analyses; (b) Description of each environmental variable included; (c) Pearson correlation coefficients between environmental variables to assess multicollinearity; (d) Identified outlier loci based on the environmental association tests; (e) Number of associated SNPs per environmental variable.

**Table S3. (a)** Ancestry proportions estimated from ADMIXTURE analyses (K = 12, Supplementary Figure S2). This table summarizes the genetic composition of individuals, revealing the proportion of ancestry assigned to each of the 12 clusters (attached as xls). **(b)** LAMP analysis results showing the estimated ancestry proportions for dingo and dog populations. The table includes details on average proportions and individual variation (attached as xls). **(c)** ELAI analysis results presenting the inferred local ancestry across populations. The table summarizes the inferred ancestry proportions at each chromosome level for dingoes and dogs (attached as xls). **(d)** GHap analysis results presenting the inferred local ancestry across populations (attached as xls).

**Table S4.** Genomic coordinates of the eight ancestry deserts detected in dingoes (defined as ≥10 consecutive SNPs with <0.1 % dog ancestry following Sankararaman et al., 2014) (attached as xls).

**Table S5.** Results of dN/dS analysis for each introgression block on chromosomes 9 and 27. The table lists genes within each block and their dN/dS ratios, indicating potential selective pressures acting on these genes (attached as xls).

**Table S6.** Genes identified within the inverted block on chromosome 9 and introgressed blocks on chromosomes 13, 14, 24 and 27, along with their known functions. This table provides insights into the potential biological roles of the introgressed genes (attached as xls).

**Table S7.** Bayesian genomic cline summary by dingo population. Hybrid index (HI) summarizes individual dog ancestry proportions within each population (mean, SD, and 5th/50th/95th percentiles). SDc and SDv are the posterior medians of the dispersion of genomic cline centers and slopes, respectively, and quantify locus-specific heterogeneity in introgression across loci.

| Population | N | HI mean | HI SD | HI 5% | HI 50% | HI 95% | SDc (median) | SDv (median) |
|---|---|---|---|---|---|---|---|---|
| **BigDesert** | **15** | **0.168** | **0.009** | **0.156** | **0.167** | **0.182** | **1.111** | **0.695** |
| **Captive** | **36** | **0.217** | **0.098** | **0.139** | **0.187** | **0.452** | **1.057** | **0.606** |
| **East** | **79** | **0.258** | **0.064** | **0.192** | **0.242** | **0.365** | **0.759** | **0.424** |
| **South** | **45** | **0.253** | **0.031** | **0.210** | **0.246** | **0.313** | **0.870** | **0.553** |
| **West** | **81** | **0.111** | **0.064** | **0.039** | **0.103** | **0.230** | **1.202** | **0.467** |
| **Central** | **88** | **0.171** | **0.071** | **0.038** | **0.200** | **0.248** | **1.261** | **0.460** |

**Table S7.** Bayesian genomic cline summary by dingo population. Hybrid index (HI) summarizes individual dog ancestry proportions within each population (mean, SD, and 5th/50th/95th percentiles). SDc and SDv are the posterior medians of the dispersion of genomic cline centers and slopes, respectively, and quantify locus-specific heterogeneity in introgression across loci.

### Supplementary Figures

**Figure S1. Admixture Analysis of Dingo and Dog Populations at Different K Values.** Results of the Admixture analysis for the combined dataset of dingoes and dogs, showing the inferred population structure at different values of K (K = 2 to K = 12). Each vertical bar represents an individual, with colors indicating the proportion of ancestry assigned to each inferred genetic cluster. The left section of each panel corresponds to dingo populations, while the right section represents dog populations. As K increases, the analysis reveals finer-scale genetic structure within both dingo and dog populations. The optimal K value, determined through cross-validation, is K = 12, indicating the presence of twelve distinct genetic clusters across the dataset. **Note: This figure is provided as a separate high-resolution file.**

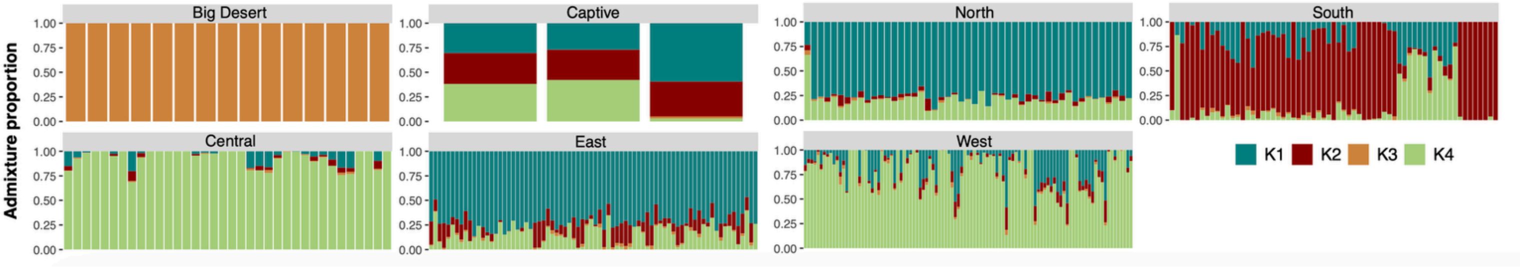


**Figure S2. Individual admixture proportions for dingoes across six geographic populations and a captive group.** Each vertical bar represents one individual, with colors indicating the proportion of ancestry assigned to each of the four genetic clusters (K1-K4). Populations are arranged by region: Big Desert (K3 only), Central, East, West, North, South, and Captive. The x-axis lists individuals within each population, and the y-axis shows the admixture proportion (0-1). Clusters K1 (teal) and K2 (maroon) predominantly characterize eastern and southern dingoes, while K3 (orange) and K4 (light green) dominate in Big Desert and central/western populations, respectively.

**(a)**

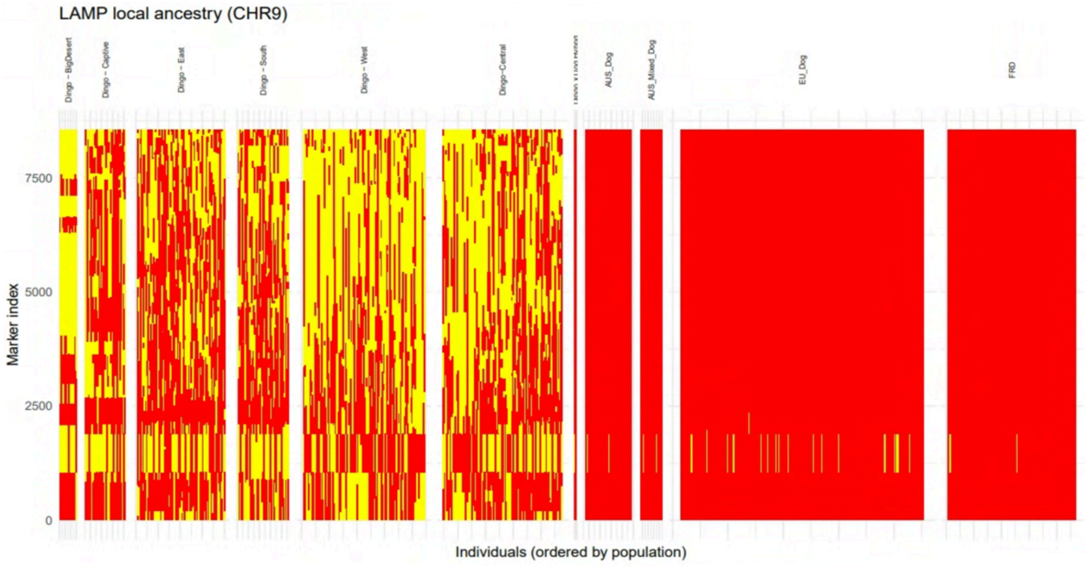


**(b)**

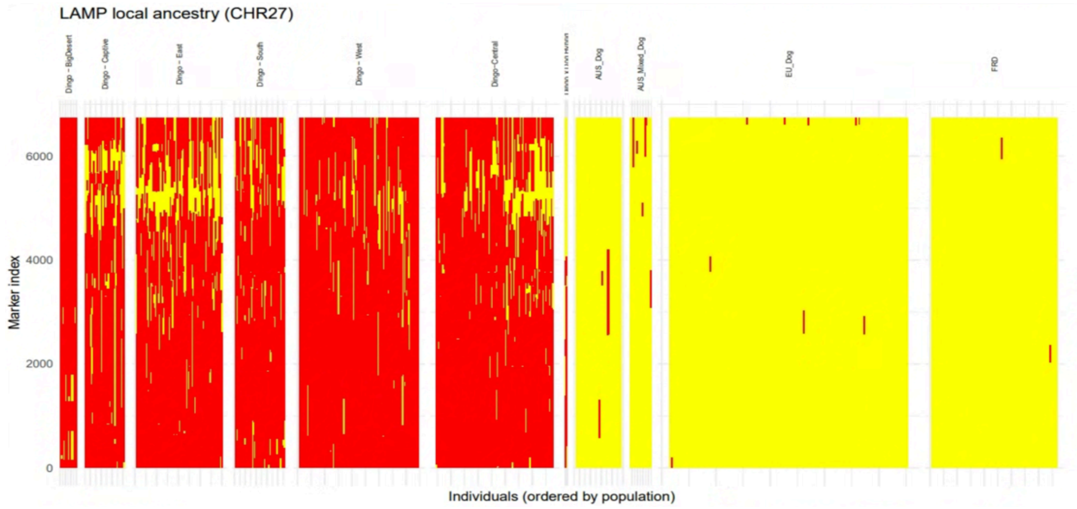


**Figure S3. Local Ancestry Plots Across Dingo and Dog Populations Using LAMP.** LAMP plots for chromosomes 9 **(a)** and 27 **(b)**, illustrating local ancestry patterns across dingo and dog populations. Each plot depicts the inferred proportion of dog ancestry along the chromosome, highlighting regions of elevated or reduced introgression. Notably, the signal observed on chromosome 9 corresponds to a known chromosomal inversion present in some domestic dogs but absent in dingoes, which was initially interpreted as a candidate region for introgression. This pattern likely reflects extended haplotype similarity due to suppressed recombination, rather than gene flow. **Note: Full sets of LAMP plots for all chromosomes (1-38) and all populations, ordered by population, are provided as supplementary material in a compressed ZIP folder.**

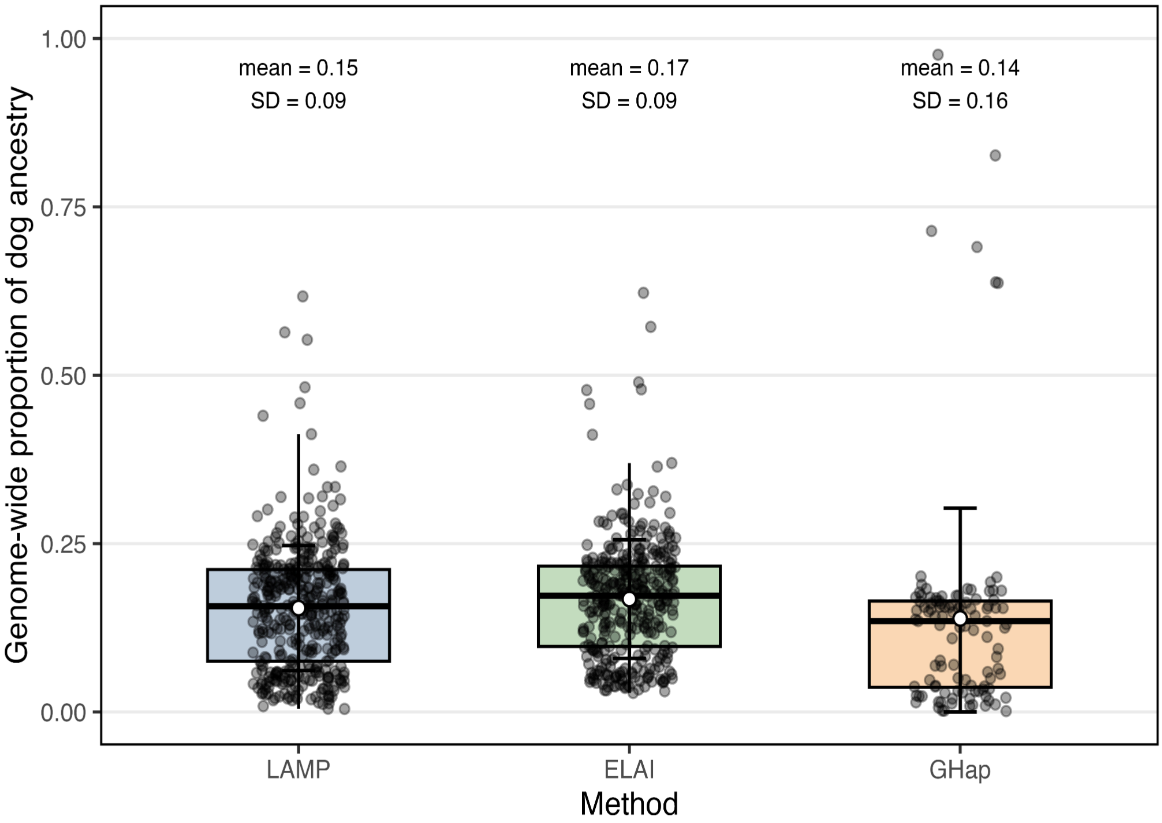


**Figure S4. Genome-wide dog ancestry estimates across local ancestry methods.** Distribution of individual-level genome-wide proportions of dog ancestry in dingoes inferred using three local ancestry-based approaches: LAMP, ELAI, and GHap. Each point represents a single individual (jittered for visibility). Boxplots show the median and interquartile range, with whiskers indicating the typical range of variation. White circles indicate mean values, and vertical bars represent ±1 standard deviation. Estimates are summarized across all dingo populations.

(a)

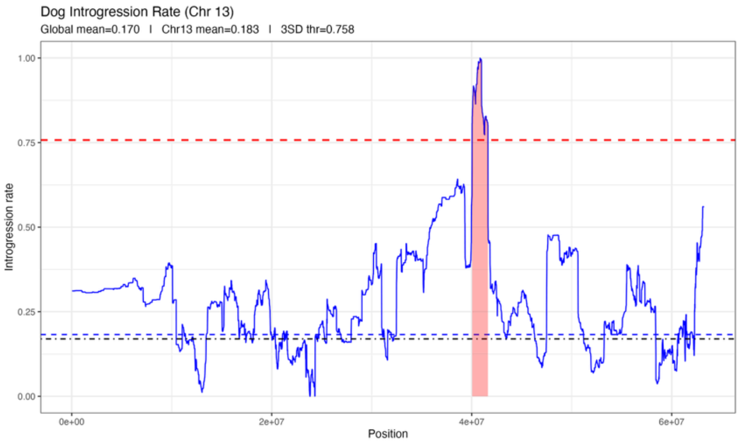
Dog Introgression Rate (Chr 13)
Global mean=0.170 | Chr13 mean=0.183 | 3SD thr=0.758
Introgression rate
Position
0.00
0.25
0.50
0.75
1.00
0e+00
2e+07
4e+07
6e+07

(b)

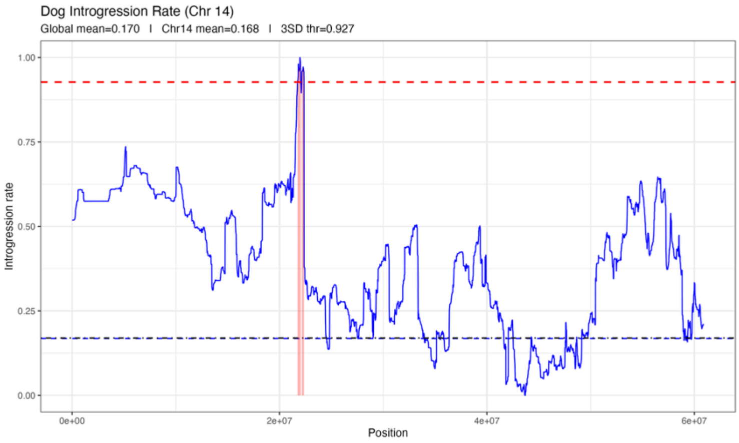
Dog Introgression Rate (Chr 14)
Global mean=0.170 | Chr14 mean=0.168 | 3SD thr=0.927
Introgression rate
Position
0.00
0.25
0.50
0.75
1.00
0e+00
2e+07
4e+07
6e+07

(c)

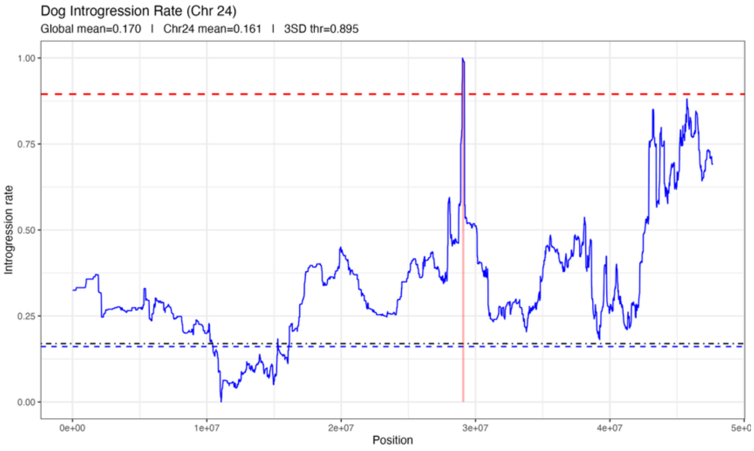
Dog Introgression Rate (Chr 24)
Global mean=0.170 | Chr24 mean=0.161 | 3SD thr=0.895
Introgression rate
Position
0.00
0.25
0.50
0.75
1.00
0e+00
1e+07
2e+07
3e+07
4e+07

**Figure S5. Dog introgression rate along chromosomes 13 (a), 14 (b) and 24 (c) based on ELAI analysis.** The x-axis is genomic position (bp) along the chromosome and the y-axis is the estimated introgression rate. Blue curves show the normalized per-SNP dog-to-dingo introgression rate (0-1) across each chromosome. The red dashed line marks the chromosome-specific detection threshold (chromosomal mean + 3 × SD of the introgression distribution), and the pale red shading highlights contiguous windows that exceed this cutoff (candidate introgressed tracts). The dashed blue line indicates the mean introgression rate on that chromosome, while the dash-dotted black line shows the genome-wide mean introgression rate. These introgression rates were calculated for the admixed dingoes only and therefore are considerably higher than population-level introgression rates.

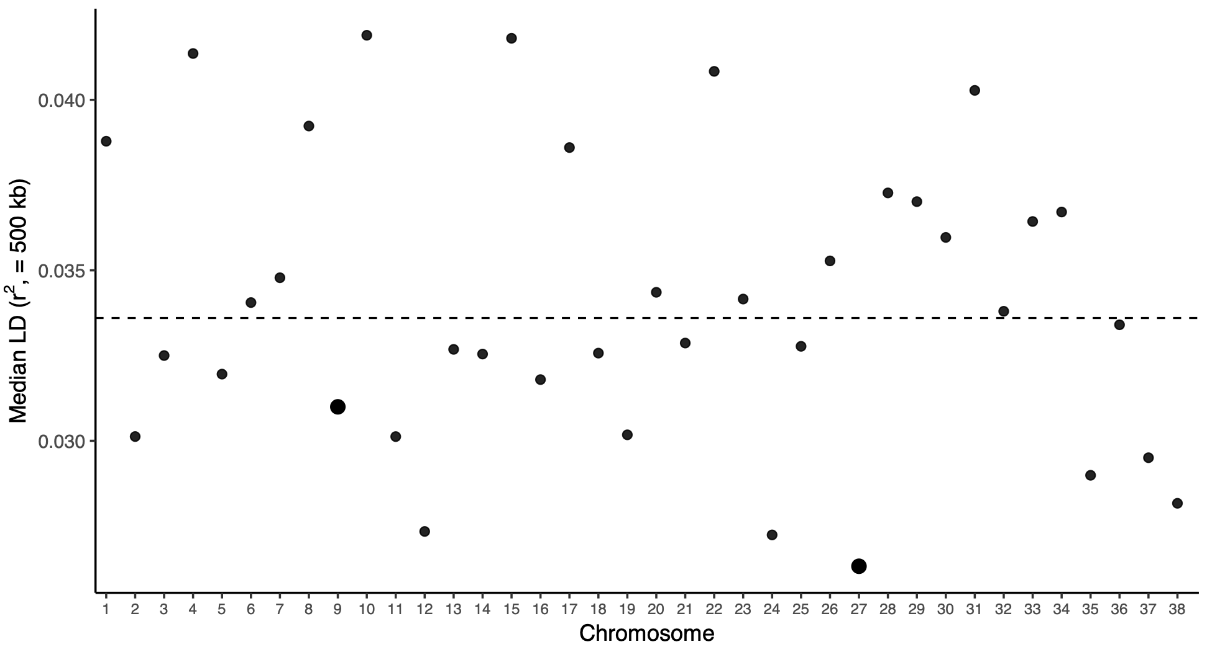


**Figure S6. Genome-wide distribution of linkage disequilibrium across autosomes.** Median linkage disequilibrium (LD; $R^2$) calculated for SNP pairs separated by up to 500 kb was summarized for each autosomal chromosome (1–38). Each point represents the chromosomal median LD value, and the dashed horizontal line indicates the genome-wide median across all autosomes.

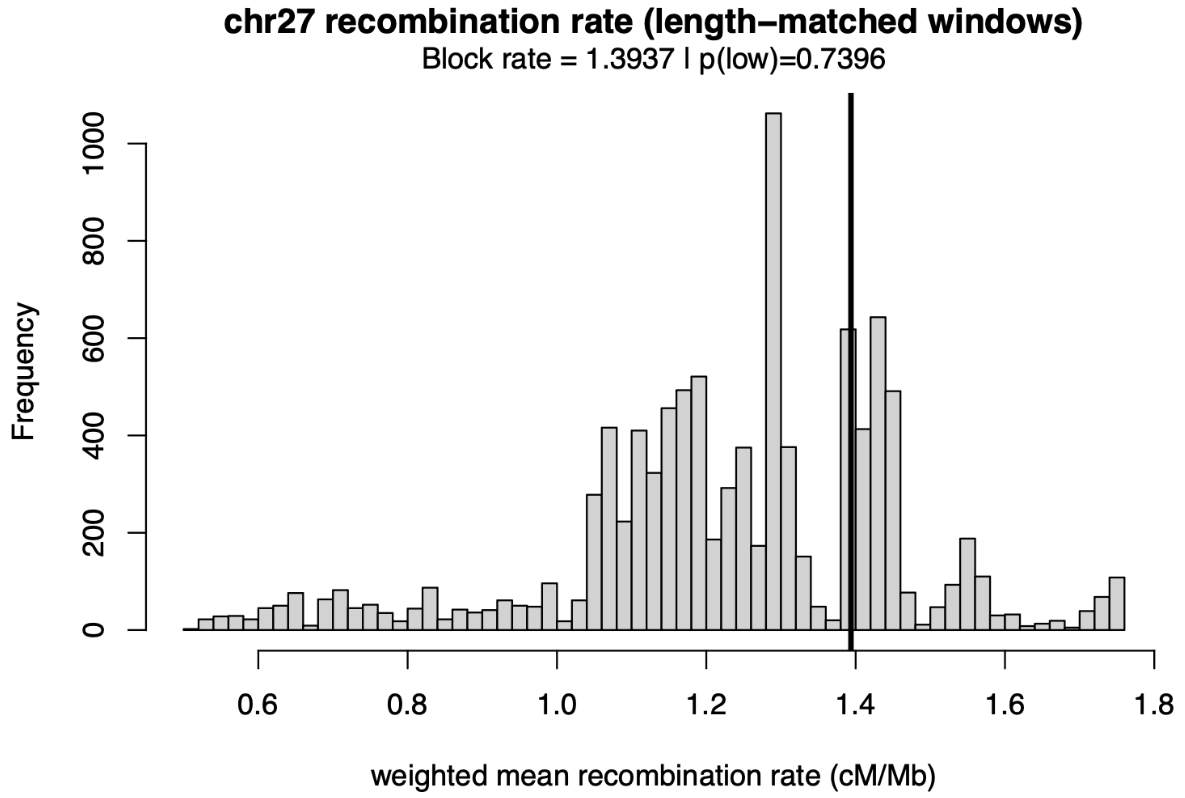


**Figure S7. Distribution of length-weighted mean recombination rates (cM/Mb) across chromosome 27** estimated from 10,000 random windows matched to the size of the chromosome 27 candidate block. The vertical line indicates the recombination rate of the focal block (1.39 cM/Mb), which falls within the chromosome-wide distribution (empirical p for low recombination = 0.74), indicating no evidence that the block overlaps a recombination coldspot.

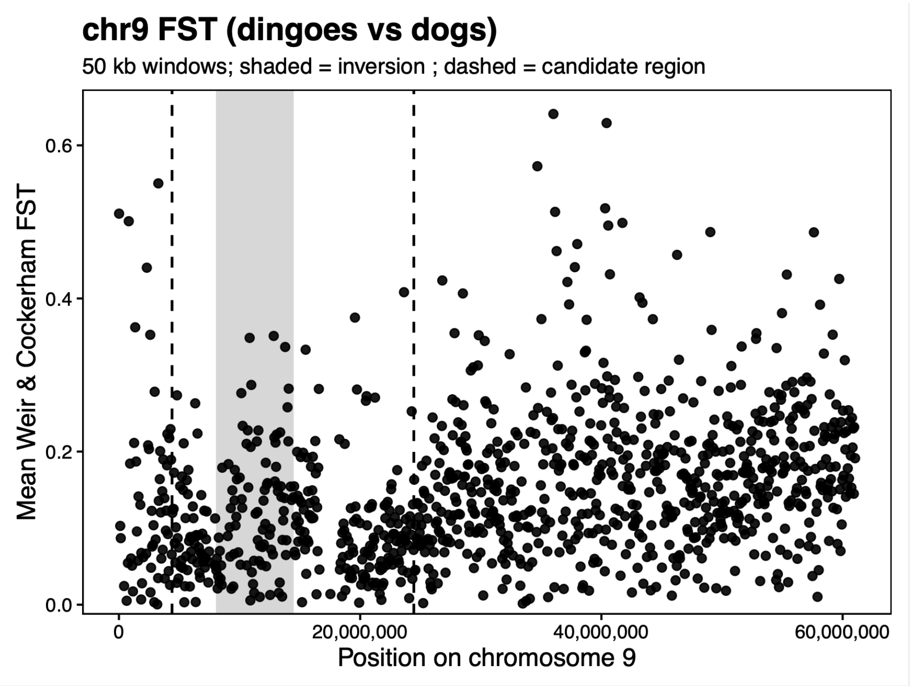


**Figure S8. Weir and Cockerham FST across chromosome 9 between dingoes and domestic dogs.** Mean FST values were calculated in 50 kb non-overlapping windows. The shaded grey area indicates the chromosomal inversion on chromosome 9 described by Field et al. (2022), while dashed vertical lines delimit the broader genomic region initially inferred as putatively introgressed.

**Figure S9. Ancestry deserts across chromosomes showing locally reduced dog ancestry.** Chromosome-specific profiles of dog introgression rates inferred from ELAI, illustrating regions of exceptionally low dog ancestry ("ancestry deserts") identified across eight autosomes. For each chromosome, the blue line shows the normalized per-SNP dog introgression rate along genomic position. The dashed horizontal line indicates the chromosome-specific threshold used to define ancestry deserts ($\leq$ 0.1% dog ancestry), and contiguous segments falling below this threshold were classified as ancestry deserts. **Note: Individual chromosome plots are provided as separate PDF files in the supplementary materials.**

**Figure S10. Phylogenetic Trees Based on Whole-Genome and Region-Specific SNP Data.** (a) Maximum likelihood phylogeny reconstructed with IQ-TREE using SNP data from the whole genome and including all individuals. The tree shows clear species-level clustering for wolves and domestic dogs, while New Guinea Singing Dogs (NGSD) do not form a reciprocally monophyletic clade distinct from dingoes, instead clustering closely with them—consistent with their shared evolutionary history. (b) Phylogeny based on the introgressed region on chromosome 27, showing clustering between dingoes and domestic dogs. All dingo populations except those from Big Desert and captive individuals possess haplotypes that group with different European dog clades, suggesting multiple independent introgression events. (c) Phylogeny of the chromosome 9 region, where some European domestic dogs cluster closely with dingoes. This pattern mirrors the local ancestry results and is consistent with a previously reported chromosomal inversion present in certain dog breeds but absent in dingoes (Field et al., 2022). The resulting phylogenetic signal likely reflects extended haplotype similarity maintained by suppressed recombination within the inversion, rather than genuine gene flow. **Note: Due to the large file sizes, full-resolution trees for all individuals are provided as PDFs in a compressed ZIP folder in the supplementary materials.**

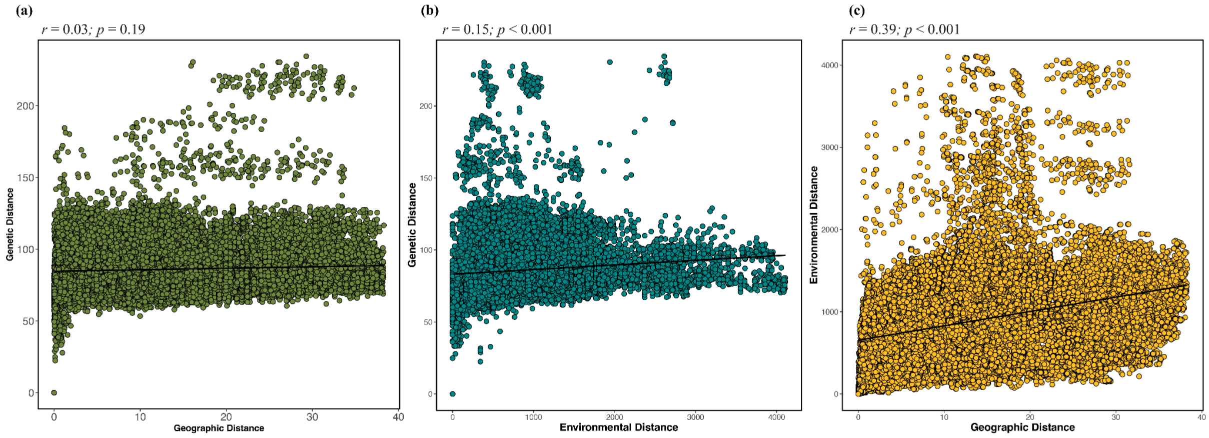


**Figure S11. Relationships Between Genetic, Geographic, and Environmental Distances.** (a) Relationship between genetic distance (y-axis) and geographic distance (x-axis). A Mantel test based on Spearman's rank correlation yielded a non-significant association between these two distances (r = 0.027, p = 0.1913). (b) Relationship between genetic distance (y-axis) and environmental distance (x-axis). The Mantel test indicated a significant positive correlation (r = 0.1515, p < 0.0001), suggesting a moderate association between genetic differentiation and environmental differences. (c) Relationship between environmental distance (y-axis) and geographic distance (x-axis). A strong and significant correlation was found (r = 0.3866, p < 0.0001), indicating a clear association between environmental variation and geographic distance.

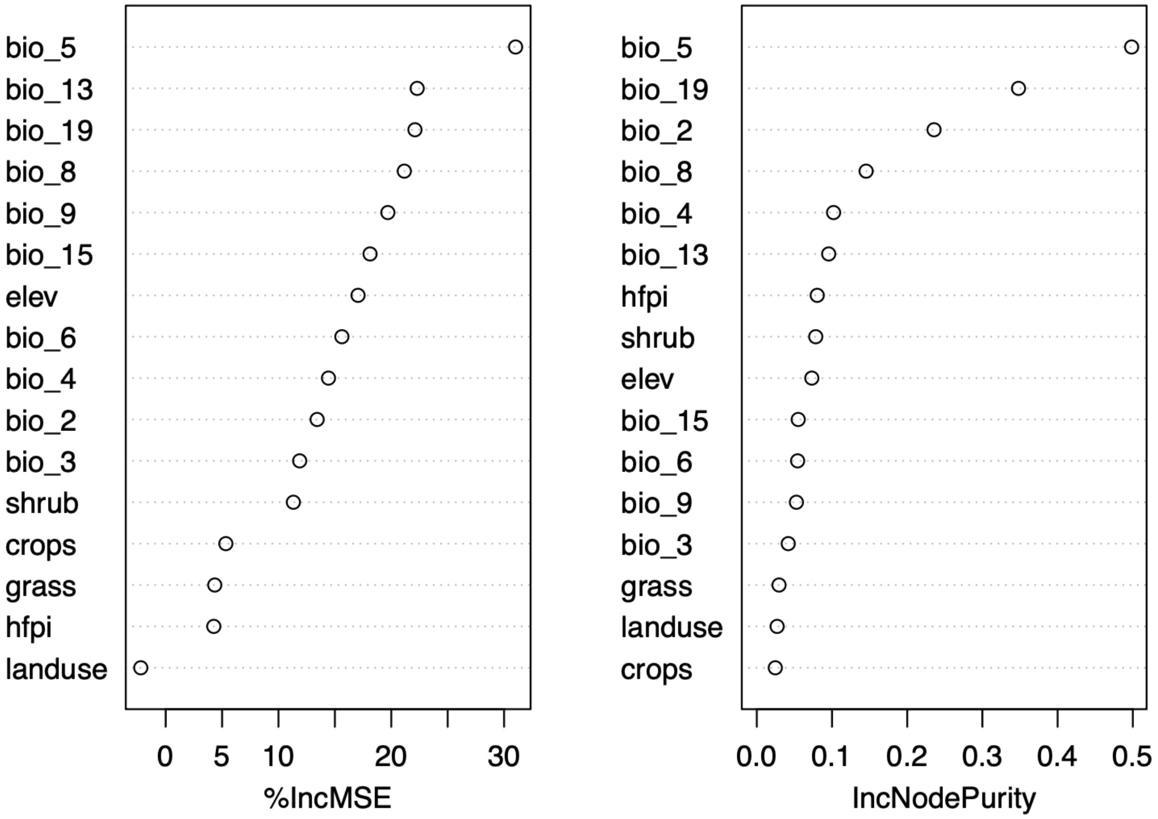


**Figure S12. Random Forest Analysis of Environmental Variables.** Variable importance from Random Forest analyses assessing the contribution of environmental and anthropogenic predictors to patterns of dog introgression in dingoes. Models were fitted using the full set of predictors described in the Methods, after removal of highly correlated variables ($|r| > 0.8$). Variable importance is shown as the percentage increase in mean squared error (%IncMSE; left panel) and the increase in node purity (IncNodePurity; right panel). Higher values indicate greater importance of a given predictor in explaining variation in introgression across individuals.
